# Metabolic exchange and energetic coupling between nutritionally stressed bacterial species, the possible role of QS molecules

**DOI:** 10.1101/2020.04.27.063917

**Authors:** David Ranava, Cassandra Backes, Ganesan Karthikeyan, Olivier Ouari, Audrey Soric, Marianne Guiral, María Luz Cárdenas, Marie Thérèse Giudici-Orticoni

**Affiliations:** CNRS, Aix-Marseille University, Bioenergetic and Protein Engineering Laboratory, Mediterranean Institute of Microbiology, 31 Chemin Joseph Aiguier, 13009 Marseille, France; Aix-Marseille University, CNRS, UMR 7273, ICR, Marseille, France; Aix-Marseille University, CNRS; Centrale Marseille, M2P2, Marseille, France

**Author notes:** DR and CB contributed equally to this work. Northwestern University, Feinberg School of Medicine, Department of Microbiology-Immunology 320 E. Superior Street Chicago, IL, 60611. MRC London Institute of Medical Sciences (LMS), Imperial College London, Hammersmith Hospital Campus, Clinical Research Building, 5th floor, Du Cane Road, London W12 0NN United Kingdom.

## Abstract

To clarify the principles controlling inter-species interactions, we previously developed a co-culture model with two anaerobic bacteria, *Clostridium acetobutylicum* and *Desulfovibrio vulgaris* Hildenborough, in which nutritional stress for *D. vulgaris* induced tight cell-cell inter-species interaction. Here we show that exchange of metabolites produced by *C. acetobutylicum* allows *D. vulgaris* to duplicate its DNA, and to be energetically viable even without its substrates. Physical interaction between *C. acetobutylicum* and *D. vulgaris* (or *Escherichia coli* and *D. vulgaris*) is linked to the quorum-sensing molecule AI-2, produced by *C*. *acetobutylicum* and *E. coli*. With nutrients *D. vulgaris* produces a small molecule that inhibits *in vitro* the AI-2 activity, and could act as an antagonist *in vivo*. Sensing of AI-2 by *D. vulgaris* could induce formation of an intercellular structure that allows directly or indirectly metabolic exchange and energetic coupling between the two bacteria.

## Introduction

Microbial communities are ubiquitous and exert a large influence in geochemical cycles and health^1–4^. In natural environments, stress factors such as nutrient deficiencies and the presence of toxic compounds can induce interactions between microorganisms from the same or different species and the establishment of communities which can occupy ecological niches otherwise inaccessible to the isolated species^5,6^. Interactions between microorganisms can affect the behavior of the community either positively or negatively^7^.

For studying ecological communities, it is crucial to understand how the different members communicate with each other and how this communication is regulated. Interactions may occur either by release of molecules into the environment^8^, or by direct contact between the microorganisms through structures like nanowires^9^ or nanotubes^10^. Dubey and Ben-Yehuda^10^ were the first to demonstrate a contact-dependent exchange of cytoplasmic molecules *via* nanotubes in *Bacillus subtilis* which contributes to proper colony formation^11,12^. The evolution of how metabolites came to be transferred between bacteria, and its functioning today, were both well described in a recent review^13^.

The type and extent of nutritional interactions between microbes partly determine the metabolism of an entire community in a given environment^14^. Very little is known about the molecular basis of interactions between species, as this is difficult to investigate, especially in Nature, on account of community complexity. The use of a synthetic microbial ecosystem has considerable interest because the reduced complexity means that the investigation is more manageable, allowing not only identification of the specific community response, but also description of the different events at the molecular and cellular level^15^.

To further investigate interactions between bacterial species we developed a synthetic microbial consortium constituted by two species: *C. acetobutylicum*, Gram-positive, and *D. vulgaris*, Gram-negative, sulfate reducing. There are both involved in anaerobic digestion of organic waste matter^16,17^. Glucose, a substrate that cannot be used by *D. vulgaris*^16^, is the sole carbon source in the consortium, and the consortium produces three times more H_2_ than *C. acetobutylicum* alone, *D. vulgaris* even grows in the absence of sulphate, its final electron acceptor for the respiration process^18^. Although *D. vulgaris* can ferment lactate, a metabolite produced by *C. acetobutylicum*, this process is greatly inhibited by high H_2_ concentrations, preventing *D. vulgaris* from growing in the absence of methanogens^19^. We observed a form of bacterial communication between adjacent cells of both types of bacteria by cell-cell interaction, in conditions of nutritional stress, with exchange in both directions of cell material, associated with the modification of the metabolism evidenced by a higher production of CO_2_ and H_2_^18^. In some cases, the interactions between *C. acetobutylicum* and *D. vulgaris*, resembled those described by Dubey and Ben-Yehuda^10^. Moreover, this type of cell-cell interactions has been seen also in other systems, which give support to its existence and functionality^10,13^.

Nutritional stress appears crucial to induce physical contact between bacteria, as this interaction was prevented by the presence of lactate and sulfate, nutrients of *D. vulgaris*. Furthermore, Pande *et al*^20^ in a synthetic co-culture of *E. coli* and *Acinetobacter baylyi*, after depletion of aminoacids such as histidine and tryptophan by genetic manipulation, observed nanotubular structures between the auxotrophs allowing cytoplasmic exchange. As in our case, the communication between the mutants was prevented by the presence of the nutrients. The formation of nanotubes between aminoacid-starved bacteria might be a strategy to survive under aminoacid limiting conditions^13^. Further evidence for the role of cell-cell connections to exchange nutrients can be found in these recent reviews^21–23^.

Altogether these studies suggest that for some species, cell-cell interaction (either by tight cell junctions, nanotube formation, vesicle chains, or flagella) can allow them to overcome nutrient starvation and that many materials, from small molecules to proteins or plasmids, can be passed from one cell to another. However, this requires not only an energetic investment to establish the connecting structures, but also to find the suitable partners. Several questions arise: what sorts of signals are involved? What is the molecular mechanism? The fact that in several cases a nutritional stress induces interaction, but addition of nutrients prevents it, raises the question whether there is a distress signal that is released from the starving bacteria, and another (a quenching factor) when nutrients are there? Specific signalling between cells is of great importance in the proper development of the community and in its stability in the long term^24^.

Here we partially answer these questions; in particular, we examine if the nutritional stress, which appears to be necessary, is also sufficient. We have investigated the possible role that quorum-sensing molecules could play in attaching the two bacterial cells involved in the consortia previously studied: *D. vulgaris* and *C. acetobutylicum* or *D. vulgaris* and *E. coli*, and we have examined how satisfactory the energetic state of *D. vulgaris* is in the co-culture when it is deprived of sulfate, and why the presence of nutrients prevents interaction between the bacteria.

## Results

### Tight bacterial interaction in the co-culture allows *D. vulgaris* to be metabolically active and to grow by using carbon metabolites produced by *C. acetobutylicum*

Our previous results demonstrated that conditions of nutritional stress of *D. vulgaris* induce a tight interaction between *D. vulgaris* and *C. acetobutylicum*, in co-culture, which allows the exchange of cytoplasmic molecules and the growth of *D. vulgaris*. If the tight interaction is prevented, *D. vulgaris* cannot grow^18^. A similar phenomenon was observed between *D. vulgaris* and *E. coli* DH10B^18^ (Supplementary Figure 5). This growth suggests an adequate energetic state of *D. vulgaris* in co-culture, despite of the lack of sulfate, its final electron acceptor, and shows that it can use metabolites from *C. acetobutylicum*, as *D. vulgaris* cannot use glucose^16^.

To evaluate the physiological impact that *C. acetobutylicum* has on *D. vulgaris* in co-culture, we labelled *D. vulgaris* cells with Redox Sensor Green (RSG), a small molecule that can easily pass through the membranes of Gram-negative and Gram-positive bacteria, used as a respiration sensor to identify metabolically active cells. It has been successfully tested on numerous bacteria, as an indicator of active respiration in pure or co-cultures^9,25^. If *D. vulgaris* became metabolically active due to its physical interaction with *C. acetobutylicum*, then a RSG fluorescence should be detected as for *D. vulgaris* cultivated in Starkey medium (lactate/sulfate respiration) (Fig. 1a). As expected, in respiration conditions *i.e*. in the presence of lactate and sulfate, substrates of *D. vulgaris, D. vulgaris* in pure culture grows well and all the cells show intense RSG fluorescence, in contrast to the culture in GY medium, where there is no growth and very few cells fluoresce (Fig. 1a and b). As a control, we added CCCP, which by dissipating the proton gradient and abolishing the ATP synthesis, tightly impacts RSG fluorescence (Supplementary Fig.1). When *D. vulgaris* is labelled with RSG, as above, and then cultivated for 20h with *C. acetobutylicum* in GY medium, despite the lack of sulfate it shows a significant RSG fluorescence indicating that the cells have enough reducing power to reduce redox green and are metabolically active (Fig. 1c-e). Furthermore, *C. acetobutylicum*, which was not labelled by RSG at the beginning, also fluoresce intensely, indicating that RSG has been transferred from *D. vulgaris*.

**Figure 1:**
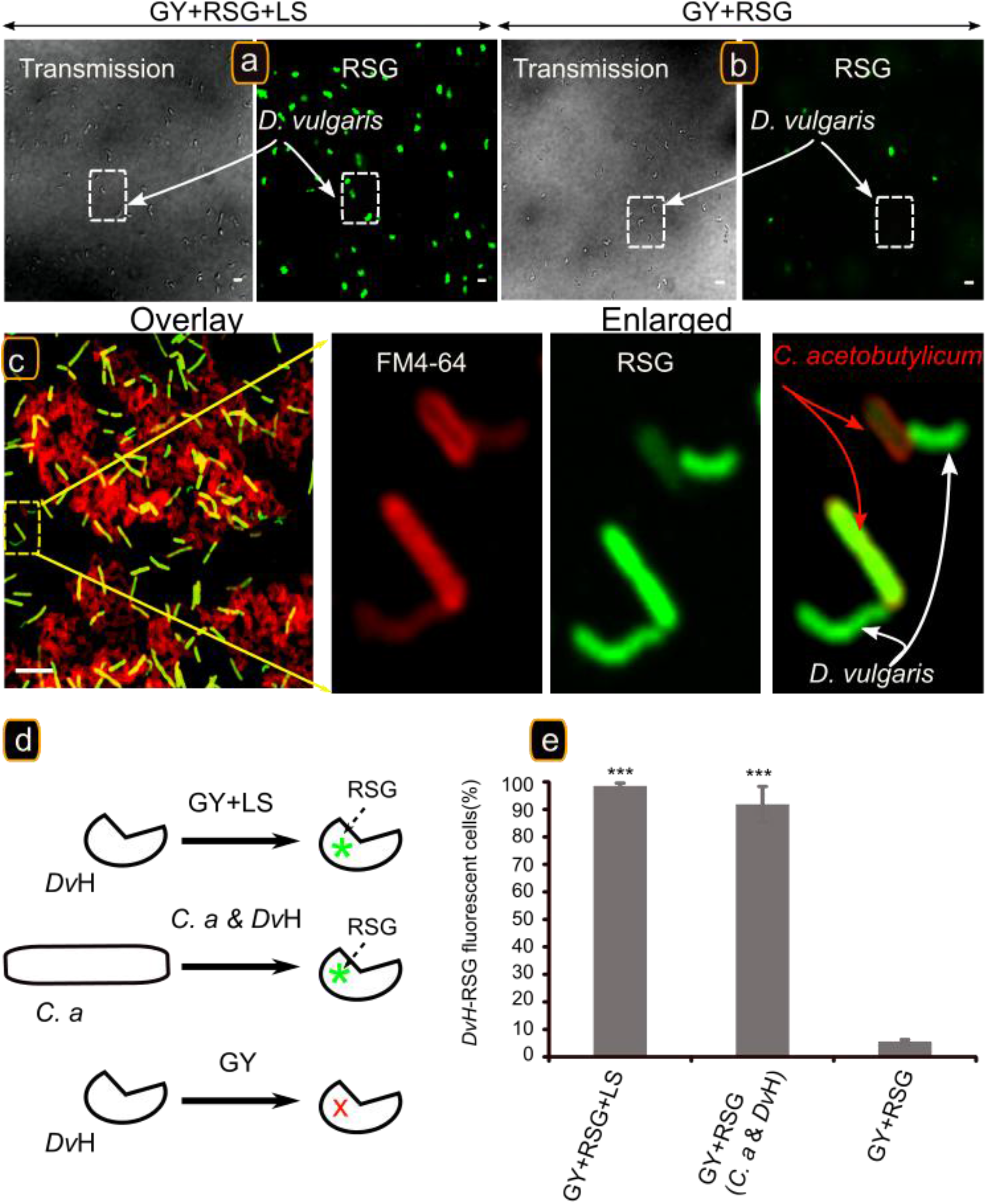
The presence of *C. acetobutylicum* is required for energetic activation of *D. vulgaris*. *D. vulgaris* growing exponentially in Starkey medium was washed twice and starved by incubation in GY medium, 20h at 37°C. A starved culture was divided into three subcultures and supplemented with 1μM of RSG (final concentration). The first was activated with 10 mM of lactate/sulfate (LS) (**a**) and the second remained starved (**b**). The two subcultures were sampled and visualized by fluorescence confocal microscopy after 20h incubation at 37°C **c)**. The third subculture was mixed with *C. acetobutylicum* cells. After 20h incubation at 37 °C, the culture was sampled, left for 5 min in contact with FM4-64 (in order to visualize the two strains) and visualized by fluorescence confocal microscopy. Scale bar, 2 μm in all panels. (n=3). (d) Schematic description of RSG treatment to identify active metabolic cells of *D. vulgaris* when the GY medium is supplemented with lactate/sulfate, in GY medium with *C*. *acetobutylicum*, and in GY medium in pure culture. Percentages of *D. vulgaris* – RSG fluorescent cells in GY medium supplemented with lactate/sulfate, in GY medium with *C. acetobutylicum*, and in GY medium in pure culture (e) Data are represented as mean ± SD with n = 3, in comparison to *D. vulgaris* in pure culture in GY medium. p-values calculated in Tukey HSD tests, * p < 0.05; ** p < 0.01; *** p < 0.001. Abbreviations: *C. a, Clostridium acetobutylicum; Dv*H, *Desulfovibrio vulgaris* Hildenborough.

To detect carbon exchange between the two bacteria, we used Stable Isotope Probing, growing *C. acetobutylicum*, either alone or in co-culture with unlabelled *D. vulgaris*, on ^13^C-glucose medium. Cells were collected at the end of the exponential phase and we separated ^13^C-DNA (heavier) from ^12^C-DNA (lighter) by density-gradient centrifugation (Supplementary Figure. 2). Analysis of the different fractions using specific gene markers for each bacterium shows that in the co-culture DNA from *D. vulgaris* is “heavy” (^13^C-labelled), indicating that metabolites derived from ^13^C-glucose were transferred between the two bacteria and used by *D. vulgaris*. Quantitative PCR emphasized the presence of *D. vulgaris* ^13^C-labelled-DNA with *C. acetobutylicum* ^13^C-labelled-DNA in the same fraction (Fig. 2). A small amount of ^12^C-unlabelled-DNA (from *D. vulgaris* and *C. acetobutylicum*) can be detected in this high-density fraction when an unlabelled co-culture is used, but negligible in relation to the total DNA. As *D. vulgaris* cannot grow on glucose or other hexoses, the ^13^C-labelled-DNA from *D. vulgaris* must have been formed using metabolites produced by *C. acetobutylicum*.

**Figure 2:**
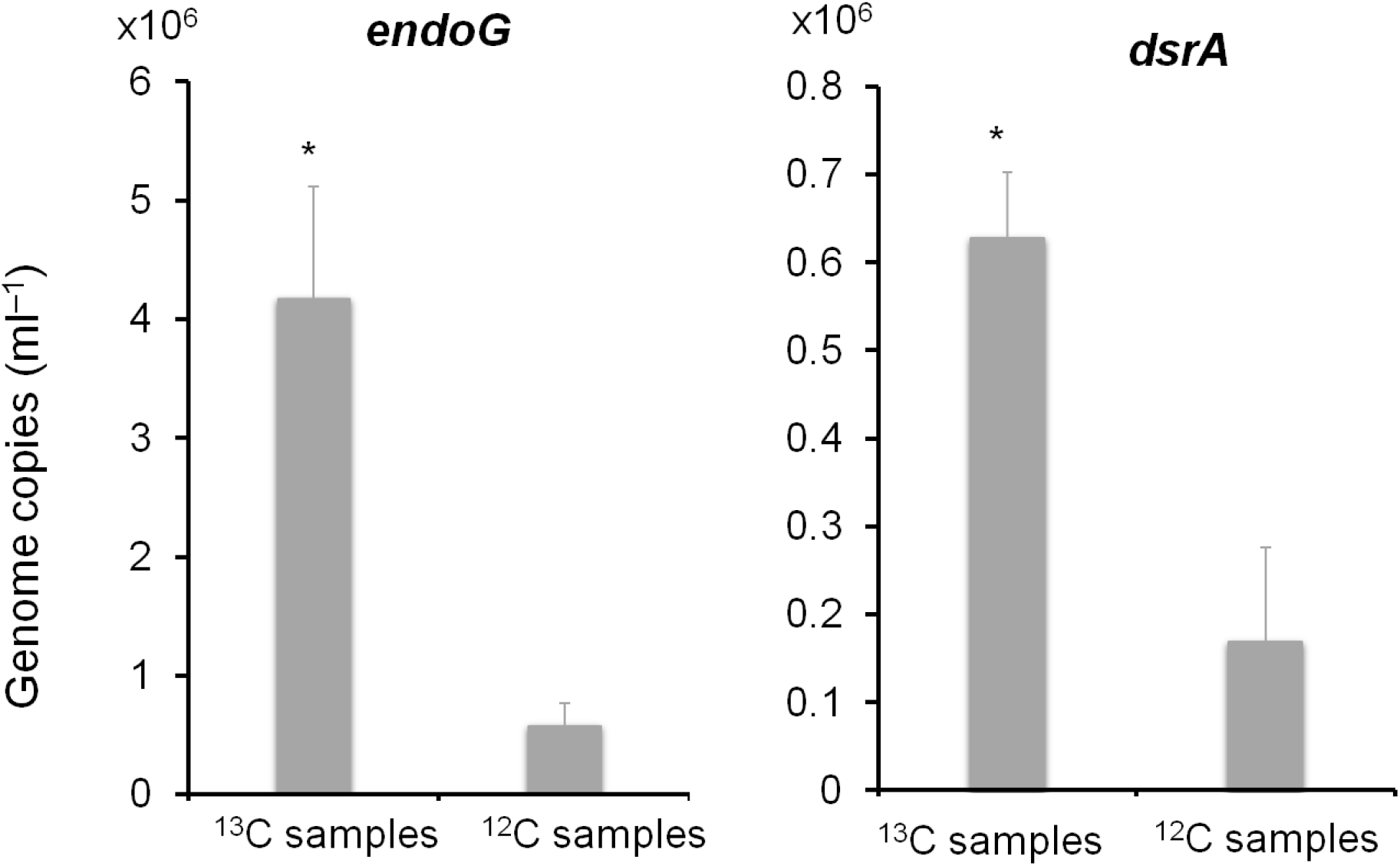
**Quantification of *Dv*H (*dsrA*) and *C*. *a* (*endoG*)** on fractions containing ^13^C from *C. a* ^13^C + *Dv*H ^12^C in ^13^C glucose medium DNA (in red) and on corresponding fractions containing ^12^C from *C. a* ^12^C + *Dv*H ^12^C in ^12^C glucose medium DNA (in green). Data are represented as mean ± SD with n = 3, in comparison to ^12^C samples. p-values calculated in Tukey HSD tests, * p < 0.05; ** p < 0.01; *** p < 0.001. Abbreviations: *C. a, Clostridium acetobutylicum; Dv*H, *Desulfovibrio vulgaris* Hildenborough

### *C. acetobutylicum* produces AI-2

As nutritional restrictions of *D. vulgaris* appear indispensable for inducing physical interactions between *C. acetobutylicum* and *D. vulgaris*, we investigated whether in addition to being necessary they were also sufficient, or if another element, such as quorum-sensing (QS) molecules, often associated with bacterial communication was required. We first looked at to homoserine lactone molecules without any success. As the co-culture is composed by an association of Gram-positive and Gram-negative bacteria, AI-2, known to be involved in interspecies communication, appears as a good candidate ^26^, and is widely accepted as the universal cell-to-cell signal in prokaryotic microorganisms^27,28^. Furthermore, although its production has not been described in *C. acetobutylicum*, it has been in *E. coli* DH10B, which can replace *C. acetobutylicum* in the co-culture^18^. *Clostridium* species are known to develop QS systems based on peptides, but QS remains relatively unknown in sulfate-reducing bacteria (SRB) although inferences on the presence of putative QS systems in them can be made^29^. However, no AI-2 signalling/sensing had been described in *C. acetobutylicum* or *D. vulgaris*. To determine whether *C. acetobutylicum* could generate AI-2-like activity, a cell-free supernatant from *C. acetobutylicum* was tested for its ability to induce luminescence in *Vibrio harveyi* BB170 AI-2 reporter strain, a classical assay for AI-2 molecule detection. This cell-free supernatant stimulated luminescence in a similar manner to cell-free supernatant from *E. coli* DH10B (AI-2 producer) (Fig. 3a).

**Figure 3:**
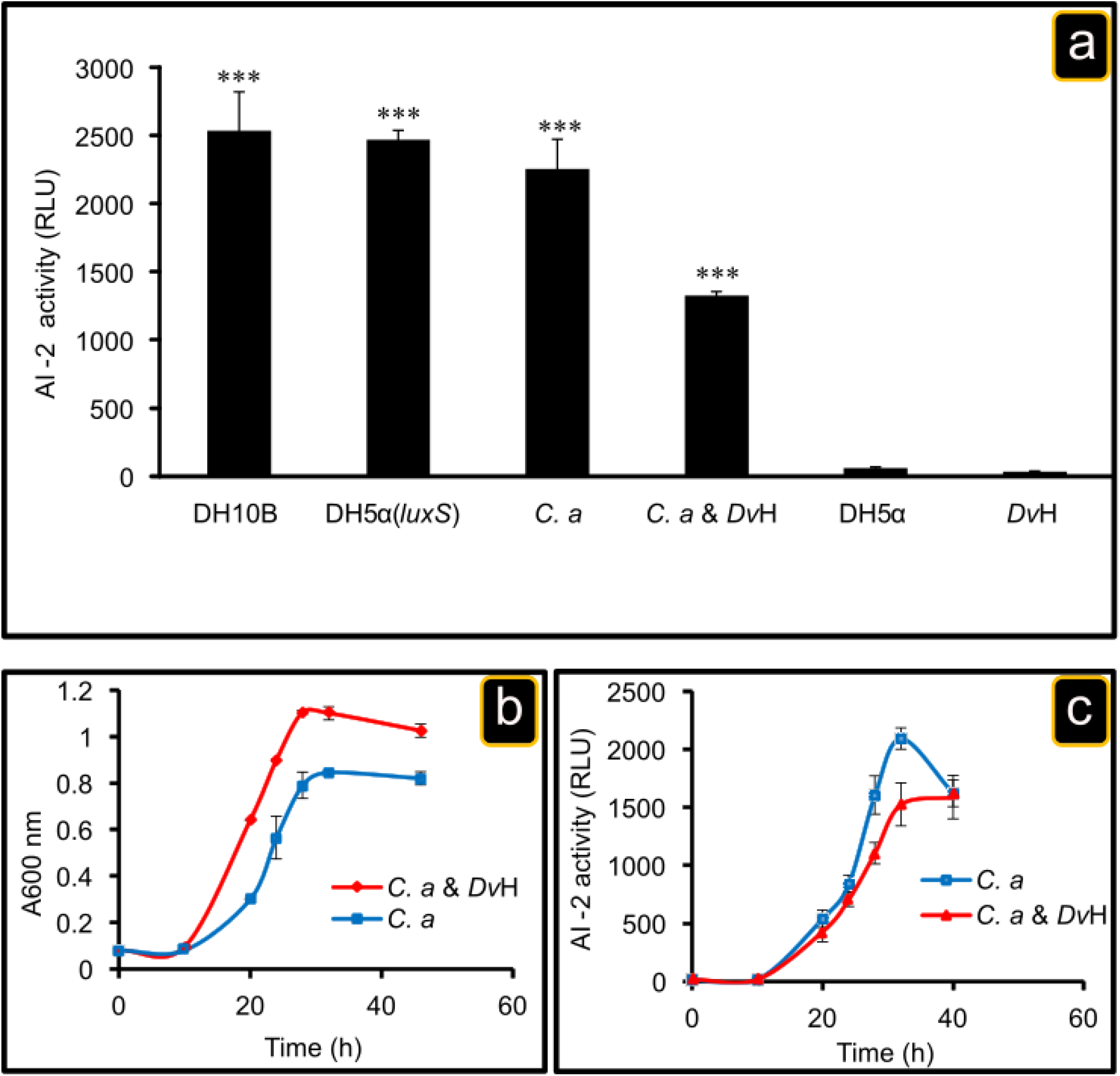
Extracellular AI-2 activity. *E. coli* DH10B (AI-2 producer; positive control), *E. coli* DH5α expressing *luxS* gene of *C. acetobutylicum, C. acetobutylicum, E. coli* DH5α (AI-2 non producer; negative control) pure cultures were grown in GY medium. *D. vulgaris* wild-type strains was grown in Starkey medium and the consortium *C. acetobutylicum* & *D. vulgaris* was grown in GY medium. The aliquots of different cultures were taken at 30h and filtered to remove cells. AI-2 activity in the cell culture supernatant was measured using the *V. harveyi* BB170 bioassay as described in Material and Methods. Data are represented as mean ± SD with n = 3, in comparison to extracellular AI-2 activity from *D. vulgaris* wild type. p-values calculated in Tukey HSD tests,*p < 0.05; **p < 0.01; ***p < 0.001 (**a**). Time course of extracellular AI-2 accumulation in pure culture of C. acetobutylicum or in co-culture *C. acetobutylicum* & *D. vulgaris*. Exponentially growing *C. acetobutylicum* in 2YTG medium and *D. vulgaris* in Starkey medium under anaerobic conditions were washed two times with fresh GY medium. Next, *C. acetobutylicum* was inoculated alone or mixed with *D. vulgari*s into GY medium at time zero and the aliquots were taken at indicated times. Cell growth was monitored by measuring the optical density at 600nm (**b**), and AI-2 activity in cell-free culture fluids was measured in a pure culture of *C. acetobutylicum* or in co-culture, using the *V. harveyi* bioluminescence assay (**c**). AI-2 activity is reported as relative light unit (RLU) of BB170 bioluminescence. Abbreviations: *C. a, Clostridium acetobutylicum; Dv*H, *Desulfovibrio vulgaris* Hildenborough ; DH5α : *Escherichia coli* DH5α; DH5α(*luxS*): *Escherichia coli* DH5α complemented *with luxS*; DH10B : Escherichia coli DH10B.

Furthermore, AI-2 activity was also detected in the co-culture *C. acetobutylicum* & *D. vulgaris*. The last step of AI-2 biosynthetic pathway is catalysed by the *luxS* gene product^30^, which is present in *C. acetobutylicum* genome (CA_C2942) annotated as S-ribosylhomocysteinase and could encode for the LuxS protein. Genetic engineering on genus *Clostridium* remains limited and requires the utilization of specific genetic tools. To avoid this, and to test if this gene is involved in AI-2 production, the putative *luxS* gene from *C. acetobutylicum* was introduced in *E. coli* DH5α (AI-2 non-producer). As hypothesised, the *E. coli* DH5α (*luxS*) expressing the *luxS* gene of *C. acetobutylicum* has AI-2 activity (Fig. 3a). In contrast, *D. vulgaris* does not have a homolog of the *luxS* gene and the cell-free culture supernatant collected from *D. vulgaris* culture, grown in Starkey medium, does not have AI-2 activity, as with the cell-free culture supernatant of *E. coli* DH5α (Fig. 3a). As *D. vulgaris* does not produce AI-2, we propose that the AI-2 molecules present in the co-culture are produced by *C. acetobutylicum*. To verify whether *C. acetobutylicum* AI-2 production follows the growth, cell-free culture supernatant from *C. acetobutylicum* or from *C. acetobutylicum* & *D. vulgaris* co-culture taken at different times were used in the *V. harveyi* bioluminescence assay, as described in Materials and Methods. As shown in Fig. 3b and 3c, *C. acetobutylicum* can synthesize functional AI-2 molecule and its synthesis follows the growth in single culture as well as in co-culture, indicating that its production is independent of the presence of *D. vulgaris*.

### Cytoplasmic exchange of molecules between bacteria in the co-culture as well as metabolic activity of *D. vulgaris* depend on the presence of AI-2

As *C. acetobutylicum* and *E. coli* DH10B, used in the previous studies^18^, both produce AI-2, this raised the question of the situation if AI-2 was not present, that is if *E. coli* DH5α, were used. So *E. coli* DH5α or *E. coli* DH10B harbouring a pRD3 plasmid containing the gene *mCherry*, was mixed with *D. vulgaris* cells lacking the *mCherry* gene but labelled with calcein, and the co-culture was analyzed by microscopy. When *D. vulgaris* cells were cultivated on GY medium with *E. coli* DH10B, more than 90% of *D. vulgaris* cells acquired a mCherry fluorescence signal after 20 h of culture (Fig. 4a panel 2, Fig. 4b and c and Supplementary Fig. 3 panel 1).

**Figure 4:**
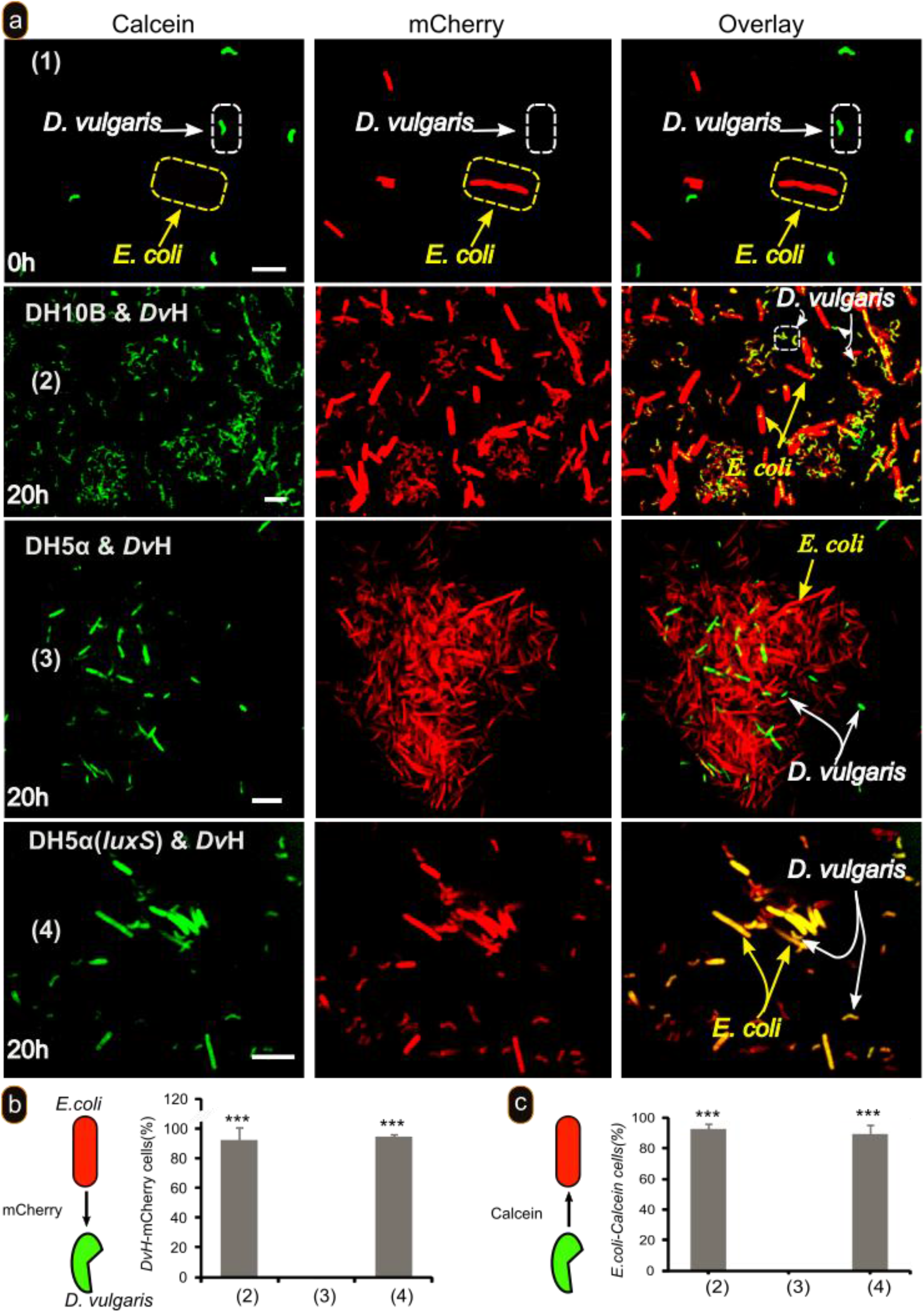
AI-2 is required for cytoplasmic molecules exchange between *D. vulgaris* and *E. coli*. *D. vulgaris* growing exponentially in Starkey medium was labelled with calcein, washed with GY medium, and mixed with *E. coli* strains DH10B (panel 2), DH5α (panel 3) and DH5α(*luxS*) (panel 4) labelled with mCherry and visualized by fluorescence confocal microscopy at time zero (panel 1) or after 20h incubation at 37°C (panels 2-4) in GY medium. Scale bar, 2 μm in all panels (a). Percentage of *D. vulgaris* labelled with mCherry in co-culture with different strains of *E. coli* (b). Percentage of *E. coli* labelled with Calcein in co-culture with *D. vulgaris* (c). Data are represented as mean ± SD with n = 3, in comparison to *E. coli* DHα (panel 3). p-values calculated in Tukey HSD tests *p < 0.05; **p < 0.01; ***p < 0.001. Abbreviations : *C. a, Clostridium acetobutylicum; Dv*H, *Desulfovibrio vulgaris* Hildenborough; DH5α : *Escherichia coli* DH5α; DH5α(*luxS*): *Escherichia coli* DH5α complemented *with luxS*; DH10B : Escherichia coli DH10B.

In contrast, no mCherry fluorescence was observed in *D. vulgaris* cells when they were cultivated with *E. coli* DH5*α* (Fig. 4 panel 3, Fig. 4b and c and Supplementary Fig. 2 panel 3). However, *D. vulgaris* cells (75%) became mCherry-fluorescent when they are co-cultured with *E. coli* DH5α expressing *luxS* gene (Fig. 4 panel 4, Fig. 4b and c and Supplementary Fig. 3 panel 2).

Taken together, these results imply that AI-2 is essential for cell-to-cell communication and exchange of cytoplasmic molecules in co-culture, but not sufficient, as nutritional stress is also required. These results may explain why in the work reported by Pande *et al*^20^ nanotubes, used to transfer amino acids, were observed between *E. coli* auxotrophe mutants, and between *E. coli* and *Acinetobacter baylyi* mutants, as in both cases there is the possibility of AI-2 produced by *E. coli*. In contrast, no nanotubes were observed between mutants of *A. baylyi* in which *luxS* is absent, according to our genome bioinformatic analysis. So when AI-2 is not produced there may be no physical interaction even if there is a nutritional stress.

In view of the necessity of AI-2 to allow growth of *D. vulgaris* in the co-culture, we tested its effect on the energetic state of the cells with RSG, as in Fig. 1. Lack of AI-2 should prevent RSG fluorescence in *D. vulgaris* by preventing physical interaction between *E. coli* and *D. vulgaris* in GY medium. Cells were incubated with RSG as described above and mixed with *E. coli* DH5α or with *E. coli* DH10B harboring the gene of *mCherry* and the co-culture was analyzed by microscopy after 20h incubation at 37 °C. *D. vulgaris* cells displayed a significant RSG fluorescence when they are mixed with *E. coli* DH10B (Fig. 5 panel 1). In contrast, no RSG fluorescence was observed in *D. vulgaris* cells in the co-culture with *E. coli* DH5α (Fig. 5 panel 2). Moreover *E. coli* DH5α does not show RSG fluorescence as *E. coli* DH10B and *C. acetobutylicum* (Fig. 1), which supports the absence of cytoplasmic exchange. In contrast, *D. vulgaris* cells co-cultivated with *E. coli* DH5α complemented with *luxS* gene displayed RSG fluorescence similar to that observed with *E. coli* DH10B (Fig. 5 panels 1 and 3). These results demonstrate that the AI-2 molecule is important for physical interaction between *D. vulgaris* and *E. coli* and thus in metabolic activation of *D. vulgaris*. All these results suggest that *D. vulgaris* can detect AI-2.

**Figure 5:**
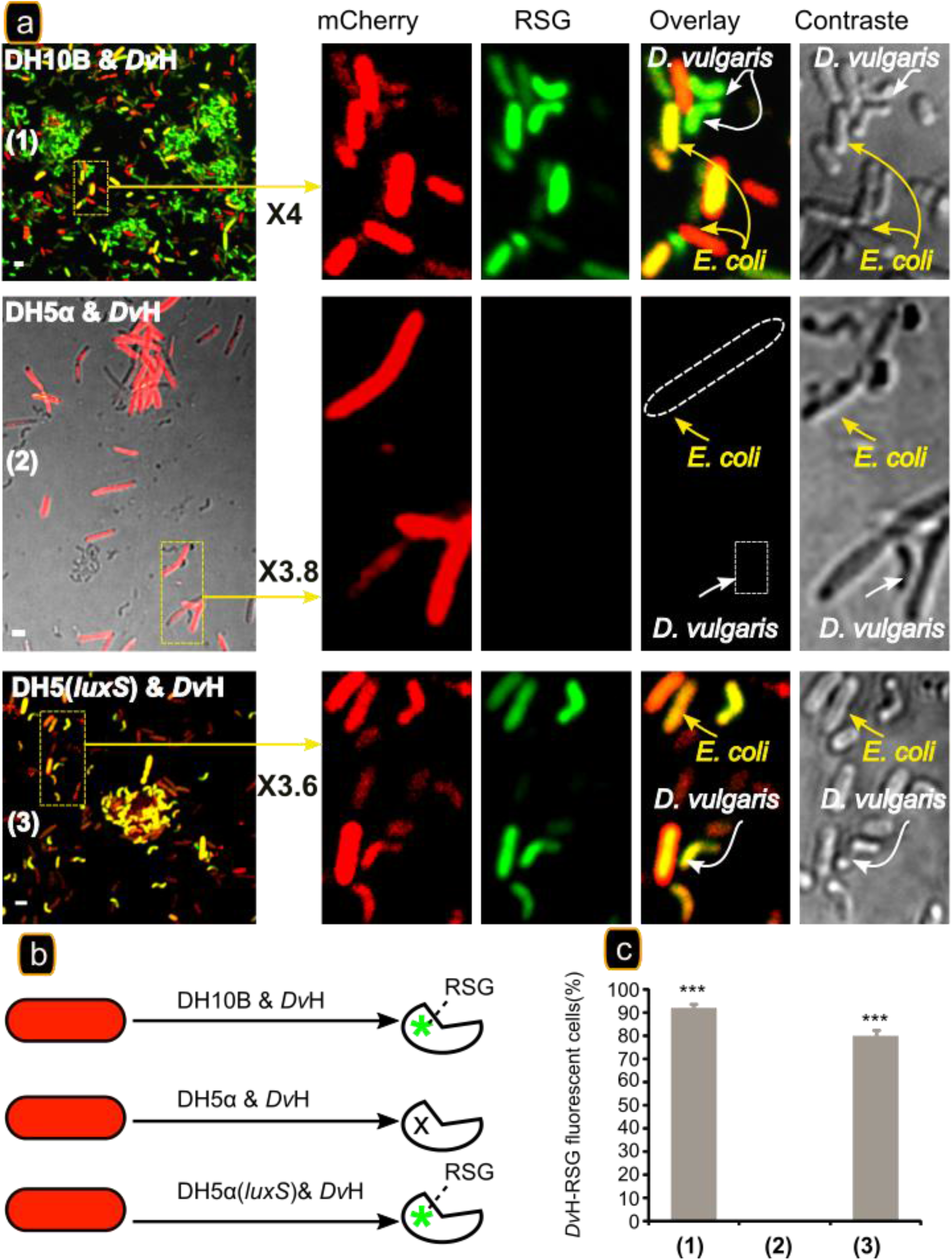
AI-2 is required for energetic activation of *D. vulgaris*. *D. vulgaris* was grown exponentially in Starkey medium was washed twice and starved by incubation in GY medium, 20h at 37°C. A starved culture was divided into three subcultures and supplemented with 1μM of RSG (final concentration). The subcultures were incubated at 37°C for 1h and then *E. coli* strains DH10B (panel 1), DH5α (panel 2) and DH5α(*luxS*) (panel 3) labelled with mCherry were added. The three subcultures were sampled and visualized by fluorescence confocal microscopy after 20h incubation at 37 °C. Scale bar, 2 μm in all panels. Schematic description of RSG treatment to identify active metabolic cells of *D. vulgaris* in co-culture with different *E. coli* strains (b). Percentages of *D. vulgaris* – RSG fluorescent cells in GY medium with *E. coli* strains (c) Data are represented as mean ± SD with n = 3, in comparison to *D. vulgaris* with *E. coli* DH5α in GY medium (non AI-2 producer). P-values calculated in Tukey HSD tests, * p < 0.05; ** p < 0.01; *** p < 0.001. *Dv*H, *Desulfovibrio vulgaris* Hildenborough; DH5α : *Escherichia coli* DH5α; DH5α(*luxS*): *Escherichia coli* DH5α complemented *with luxS*; DH10B : Escherichia coli DH10B.

### *D. vulgaris* produces an antagonist of AI-2 in the presence of sulfate and under respiratory conditions

The effect of AI-2 in the co-culture suggests that *D. vulgaris* can detect it. However, lactate and sulfate in the co-culture medium prevent physical contact between the two bacteria and transfer of cytoplasmic molecules, despite the fact that *C. acetobutylicum* and *E. coli* can produce AI-2. This suggests a regulatory mechanism linked to the presence of lactate and sulfate in the culture medium and/or to the sulfate respiration metabolism of *D. vulgaris*.

Various hypotheses may explain this: (i) in the presence of lactate and sulfate, *C. acetobutylicum* does not produce AI-2; (ii) AI-2 could be used as carbon source by *D. vulgaris;* (iii) *D. vulgaris* in the presence of sulfate produces an antagonist of AI-2.

The addition of lactate and sulfate to pure culture of *C. acetobutylicum* does not modify the production of AI-2. (Fig. 6a). In contrast, in co-culture with *D. vulgaris*, the AI-2 activity greatly decreased, even at 5 mM lactate and sulfate, and was not detected by growing the co-culture in 10 mM or higher concentrations (Fig. 6a), suggesting that in sulfate respiratory conditions *D. vulgaris* could produce one or more metabolites that inhibit the activity of AI-2. An important point is that in these conditions butyrate was produced, indicating that *C. acetobutylicum* is metabolically active and that the lack of AI-2 is not due to a metabolic inactivity of *C. acetobutylicum*.

**Figure 6:**
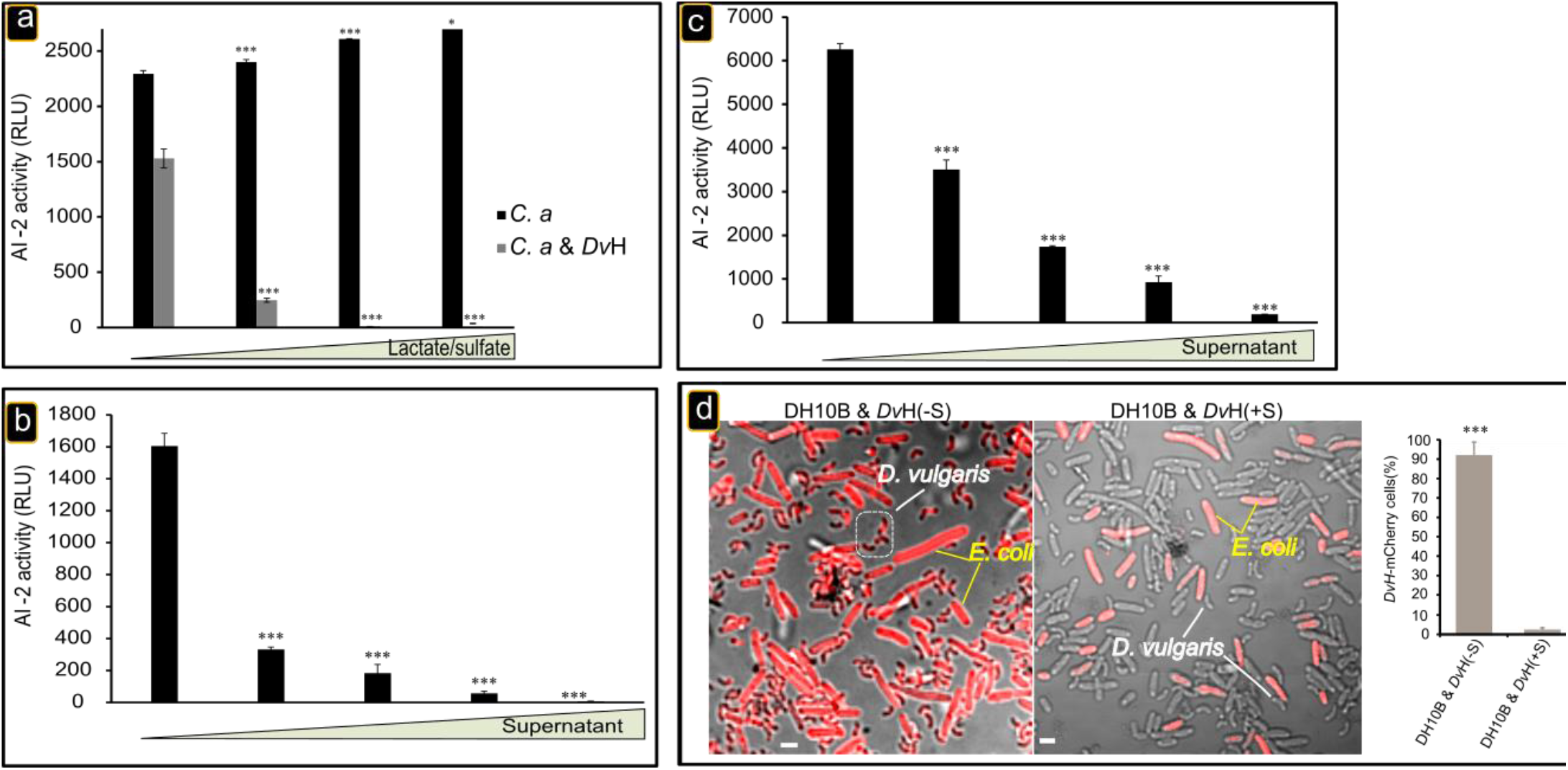
Inhibition of AI-2 activity. Addition of lactate and sulfate to the co-culture impaired AI-2 activity. Exponentially growing *C. acetobutylicum* in 2YTG medium or *D. vulgaris* in Starkey medium under anaerobic conditions was washed two times with fresh GY medium. Next, *C. acetobutylicum* was inoculated alone (black) or mixed with *D. vulgaris* (grey) into GY medium supplemented with 5, 10, and 20mM of lactate/sulfate. After 30h of incubation at 37°C, the AI-2 activity were then analyzed using the *V. harveyi* reporter strain BB170 (a). *D. vulgaris* supernatant inhibits AI-2 activity. *C. acetobutylicum (C. a*) strain was grown in GY medium during 30h at 37°C. The activity of AI-2 in the filtered samples was then analyzed using *V. harveyi* reporter strain BB170 in the presence of various quantities (1, 2, 4 and 8μL) of *D. vulgaris* filtered (0.2μm) supernatant grown on Starkey medium. AI-2 activity is reported as relative light unit (RLU) of BB170 bioluminescence (b). *E. coli* DH10B strain was grown in GY medium during 30h at 37°C. The activity of AI-2 in the filtered samples was then analyzed using *V. harveyi* reporter strain BB170 in the presence of various quantities (1, 2, 4, and 8μL) of *D. vulgaris* filtered (0.2μm) supernatant grown on Starkey medium (c). Data are represented as mean ± SD with n = 3, AI-2 activity measured for *C. acetobutylicum* in GY medium with 5, 10, and 20mM of lactate/sulfate in comparison to *C. acetobutylicum* in GY medium (black) and *C. acetobutylicum* & *D. vulgaris* in GY medium with 5, 10, and 20mM of lactate/sulfate in comparison to *C. acetobutylicum* & *D. vulgaris* in GY medium (grey) (a). In comparison to AI-2 activity measured for AI-2 produced by *C. acetobutylicum* and *E. coli* without *D. vulgaris* Starkey supernatant (b and C respectively). p-values calculated in Tukey HSD tests, * p < 0.05; ** p < 0.01; *** p < 0.001. Abbreviations : *C. a*, *Clostridium acetobutylicum*; *Dv*H, *Desulfovibrio vulgaris* Hildenborough. **d) Impact of supernatant of *D. vulgaris* growing in Starkey on cytoplasmic molecule exchange**. *D. vulgaris* growing exponentially in Starkey medium was washed with GY medium and mixed with *E. coli* DH10B (AI-2 producer) labelled with mCherry and grown in 5ml of GY medium supplemented with 100μl of *D. vulgaris* filtered (0.2μm) supernatant grown on Starkey medium. The culture was visualized by fluorescence confocal microscopy after 20h incubation at 37 °C. Scale bar, 2 μm in all panels. Data are represented as mean ± SD with n = 3, in comparison to the percentage of *D. vulgaris* fluorescent with *E. coli* DH10B in GY medium supplemented with *D. vulgaris* Starkey supernatant. p-values calculated in Tukey HSD tests, * p < 0.05; ** p < 0.01; *** p < 0.001. Abbreviations: *Dv*H, *Desulfovibrio vulgaris* Hildenborough; DH10B : *Escherichia coli* DH10B

To test the presence of molecules that could interfere with AI-2 activity in sulfate respiratory conditions, we followed the AI-2 activity present in the supernatants of *E. coli* or *C. acetobutylicum* in the presence of increasing amounts of supernatant culture of *D. vulgaris* grown in Starkey medium. Sterile filtered culture supernatant of *D. vulgaris* inhibits AI-2 activity of *E. coli* DH10B and of *C. acetobutylicum* cell-free supernatant (Fig. 6b and c), in a dose-dependent manner, indicating that *D. vulgaris*, in the presence of lactate and sulfate and independently of the presence of the other bacteria, released into the culture medium an AI-2 inhibiting compound (or a mixture of such compounds).

We followed the production kinetics of the AI-2 inhibiting compound by *D. vulgaris* by tacking samples of sterile filtered supernatant at different times of the culture of *D. vulgaris* in Starkey medium. An AI-2 inhibitor is detected 3 hours after the beginning of growth (Supplementary Fig. 4). The production kinetics suggests a QS controlled expression. Microscopy analysis of *E. coli* and *D. vulgaris* co-culture in the presence of the *D. vulgaris* supernatant show the loss of the interaction (Fig. 6d). The sulfate respiration process is probably therefore associated with the production of an antagonist or antagonists that inhibit the AI-2 activity. This production requires to the presence of sulfate, and is independent of the presence of *C. acetobutylicum* or *E. coli*.

To identify the compounds that interfere with the activity of AI-2, cell-free supernatants of *D. vulgaris*, grown in Starkey medium for 30h, were analyzed by HPLC. Different peaks (P1-P5) were recorded and their ability to inhibit AI-2 activity was determined on the supernatant of *E. coli* DH10B or on the *in vitro* synthesized AI-2 (Fig. 7a and b). Only the peak (P5) containing a 186-Da molecule (Fig. 7a et 7b, indicated by black arrow) inhibits the activity of AI-2 significantly in a dose-dependent manner (Fig. 7c). Note that the inhibitor has a molecular mass equivalent to that of AI-2, suggesting a similar type of molecule that could act as a competitive inhibitor. Different strategies for obtaining the structure did not succeed, probably because as with AI-2, there is an equilibrium between various forms^31^.

**Figure 7:**
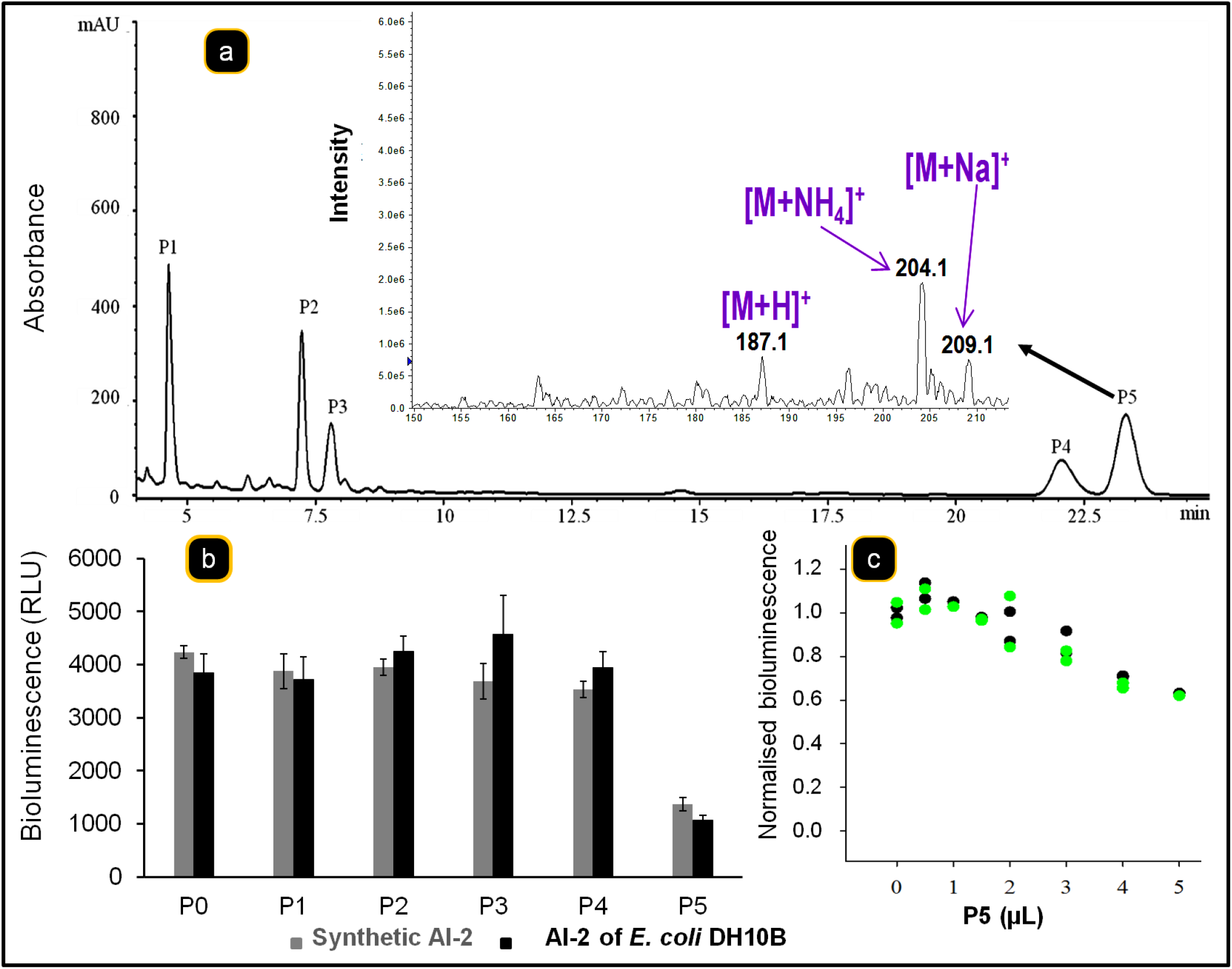
Identification of AI-2 inhibitor in *D. vulgaris* supernatant. *D. vulgaris* strain was grown in Starkey medium during 30h at 37°C and the filtered supernatant was analyzed by RP-HPLC. The peak P5 which presents an AI-2 inhibitor activity was analyzed mass spectrometry (**A**, indicated by black arrow) (a). The AI-2 inhibitor activity of different peaks collected (P1 to P5) was determined on synthetic AI-2 ((S)-4,5-Dihydroxy-2,3-pentandione, DPD) (2,5μM) or on AI-2 produced by *E. coli* strain DHI0B grown on GY medium (b). To test the hypothesis that the P5 can compete with AI-2, *V. harveyi* reporter strain BB170 was grown in AB medium (100μl) supplemented with 500nM of synthetic AI-2 and 5μL of P5 and Bioluminescence was measured each hour over the course of 4h (black circle) or of 5h (green circle) (c).

## Discussion

The established bacterial community constituted of *C. acetobutylicum* and *D. vulgaris*, or *E. coli* DH10B and *D. vulgaris*, in batch culture, and in conditions of nutritional stress for *D. vulgaris*: (i) exchanges metabolites, allowing a satisfactory energetic state and growth of *D. vulgaris*, and (ii) is regulated by AI-2 molecules that allow physical and metabolic interactions in the co-culture. Furthermore, in the presence of sulfate, *D. vulgaris* is able to produce an AI-2 antagonist. Production of the antagonist is independent of the presence of *C. acetobutylicum* or *E. coli* and may prevent formation of other consortia. Altogether, our studies revealed how QS molecules coordinate interactions between species and this modulation follows the environmental stress. Moreover, they illustrate how experiments with multiple species or synthetic ecological models can provide new insight into bacterial sociability.

The presence of *C. acetobutylicum* or *E. coli* (DH10B) allows *D. vulgaris* to be energetically viable despite being without nutrients. The observation that without *C. acetobutylicum* or *E. coli* DH10B, *D. vulgaris* is not energized explained the fact that the presence of one or the other of these two bacteria is indispensable for growth of *D. vulgaris*, as in their presence the formation of nanotube-like structures allows transfer of metabolites from one bacterium to the other.

The presence of ^13^C-DNA from *D. vulgaris* in conditions of co-culture with *C. acetobutylicum* using only ^13^C-glucose indicates that *D. vulgaris* can grow on the metabolites produced by *C. acetobutylicum* and derived from ^13^C-glucose if the two types of bacteria are in tight contact. Although *D. vulgaris* might just be using the metabolites produced by *C. acetobutylicum* and excreted to the culture medium, this is not supported by the observations that intercellular connections appear indispensable to allow growth of *D. vulgaris*^18^ as the presence of dialysis membrane that prevents the contact also prevents the growth. As in the co-culture, in absence of sulfate (electron acceptor) and in conditions of tight contact between *D. vulgaris* and *C. acetobutylicum*, (i) *D. vulgaris* uses the metabolites produced by *C.acetobutylicum*, to grow; (ii) *D.vulgaris* cannot grow on lactate fermentation because of inhibition by H_2_; (iii) growth requires the existence of a respiratory metabolism. Thus *C. acetobutylicum* may be acting as final electron acceptor through a mechanism not yet elucidated.

How these kinds of interactions are initiated and controlled is at present poorly understood. Ben-Yehuda’s group showed that YmdB is involved in the late adaptive responses of *B. subtilis* in the early stage of nanotube development^12^. In various types of cells, it is the cell undergoing stress that develops nanotube formation, suggesting that this might be directly induced by stress and constitutes a defense mechanism^32^ as apparently also in this consortium.

Surprisingly, the role and the consequence of the QS molecules, well described in pure culture, are poorly investigated and understood in bacterial consortia, closer to those found in Nature. Recently one study investigates this question using mathematical model to demonstrate/ propose how QS control the population trajectories in synthetic consortium^33^. However, this has attracted the attention of researchers studying mixed cultures in bioreactors for treating waste water. Although the real mechanism involved in QS regulation, of complex microbial consortia, remained to be elucidated, studies of this type have shed light on it^34,35^.

The QS molecule AI-2 is crucial for metabolic interaction between the two bacteria of the co-culture, as its absence prevents the metabolic exchange. Thus *C. acetobutylicum* produces a molecule with AI-2 activity, which had not been described before; furthermore, the *C. acetobutylicum* gene *luxS* can restore the AI-2 production of an *E coli* DH5α. Our results explain why *E. coli* can connect to other bacterial cells to exchange cytoplasmic molecules as described by Pande *et al*^21^ but not *A. baylyi* which lacks the *luxS* gene, according to our genomic analysis.

Some organisms only produce AI-2 whereas others only sense the signal^36,37^. *D. vulgaris* appears to be in this last category, as it lacks the *luxS* gene, but appears to sense AI-2. This suggests the presence of AI-2 receptors in *D. vulgaris*. However, no genes similar to *lsrB* (coding for LsrB, protein receptor of AI-2 in enterobacteria) or to *luxP* (gene coding for LuxP, the protein receptor of AI-2 in *vibrionaceae*) are present in the genome of *D. vulgaris*^16^. However, some bacteria can respond to exogenous AI-2 signal^38,39^ despite not having the genes *luxS, lsrB or luxP*. Furthermore, two proteins can bind AI-2 in *Helicobacter pylori*^40^, AibA and AibB; deletion of their genes (*hppA* and *modA*) induces deficiencies in chemotaxis and biofilm organization. By bioinformatic analysis we have detected their homologs (*oppA* and *modA* respectively) in *D. vulgaris*, and experiments are in progress to study them. So, the issue of the cellular receptor of AI-2 remains an open problem and our results may contribute to clarify it.

Although the original QS concept was focused on the detection of cell density for the regulation of gene expression, studies in microbial ecology suggest a wider function. For example, the efficient-sensing concept^41^ assumes that the ecologically relevant function of AI-2 sensing is to pre-assess the efficiency of producing extracellular effectors or “public goods”. Cooperative genes regulated by QS molecules can also be sensitive to nutrient conditions, suggesting that metabolic information is integrated into the decision to cooperate. AI-2 molecules are involved in the mechanism stimulating viable but no cultivable cell exits from dormancy, perhaps signalling to dormant cells when conditions are now favorable for growth^42,43^. This supports the idea that AI-2-dependent signalling reflects the metabolic state of the cell, and can function as a proxy for the production of effectors such as enzymes, or the formation of nanotubes. Integrating metabolic information with QS offers a possible mechanism to prevent cheating, as cells can only cooperate when they have the appropriate nutritional resources to do so, reducing the cost of cooperation to the individual cell^44^.

We demonstrate the quenching of the AI-2 activity by a QQ molecule produced by D. vulgaris in the presence of lactate/sulfate. Quorum quenching (QQ) has been suggested to be achieved in three ways: (i) blocking synthesis of autoinducers; (ii) interfering with signal receptors; and (iii) degrading the autoinducers^45–48^. As we show competition between AI-2 present in *C. acetobutylicum* or *E. coli* supernatant, or even synthetic AI-2 and an AI-2 quencher, a small molecule presents in the *D. vulgaris* supernatant, we can exclude the first and the third hypotheses. Only a few AI-2 interfering mechanisms have been reported and most of them include synthetic molecules as quencher^49,50^. Roy *et al*^31^ proposed that the antagonist (C1 alkyl analogues of AI-2) could compete with AI-2 for the LsrR transcriptional regulator in the lsr system^31,51^ and the presence of the competitor is linked to a decrease in AI-2 production.

As AI-2 consists of a group of molecules in equilibrium, not a unique defined structure^31^, analogy with enzymes and alternative substrates suggests that different types of AI-2 molecules may interact with a receptor, with only some of them inducing a response. We cannot discard the possibility that the QQ molecule identified in *D. vulgaris* supernatant could bind to an AI-2 receptor in *C. acetobutylicum* and induce an effect at the level of gene transcription that could be translated into metabolic modification. Moreover, we also cannot discard the possibility that the QQ molecule identified represents a QS signal for *D. vulgaris*.

Our analysis provides new insights into metabolic prudence^47,52^ and bacterial communication, and about how metabolic signals influence social behaviour but many details of its molecular implementation remain to be discovered. Which proteins detect the metabolic signals? How do they interact with QS regulation at the molecular level? However, one should also be cautious in using the word “signalling” because every change in a living organism affects every other, and thus acts as a signal of some kind^53,54^. In all of these studies it is important to keep in mind the ecological context, but the analysis of how the components of an ecological system influence one another has barely begun^55^. Anyway, we can see in these microbial communities established, thanks to QS molecules, the preliminary steps in the evolutionary pathway of multicellular organisms and eukaryotes.

## Materials and Methods

### Media and growth conditions

Strains were grown to steady state in Hungate tubes under anaerobic conditions, in LB medium for *E. coli* DH10B and DH5α, in Starkey medium (containing lactate and sulfate) for *D. vulgaris*^56^ and 2YTG medium for *C. acetobutylicum*^57^. The growth medium (Glucose-Yeast extract (GY) medium) used for studying the consortium was prepared with glucose (14 mM), 0.1% yeast extract, and supplemented with the similar inorganic nutrients used for the Starkey preparation (but with MgCl_2_ instead of MgSO_4_). GY medium was inoculated with either washed *D. vulgaris* or *C. acetobutylicum* or *E. coli* or with the combination of different strains to constitute an artificial consortium in a 1:1 ratio according to the absorbance at 600 nm. In some cases, the growth medium was supplemented with 5 or 10mM lactate or/and 5 or 10 mM sulfate. The experiments were carried out at least in triplicate.

### Construction of *E. coli* DH5α (*luxS*) strain

The *luxS* ORF (corresponding to gene *CA_C2942*) was amplified using the genome of *C. acetobutylicum* as a template and oligonucleotide primers *LuxS* 5’ GAAACCGGTAAAACAAAGGAGGACGTTTATGGAAAAAATCGCAAGTTTTACTG-3’ *LuxS-RevpB* 5’-GATCGATGGTACCTTATCAGTGGTGGTGGTGGTGGTGCTCTGGATAATTTAATCTATCTTCAGATATG-3’. The PCR amplification of *luxs* was digested and introduced into the AgeI and KpnI sites of pBGF4 plasmid under the control of the hydrogenase constitutive strong promoter^58^ to obtain pRD4 plasmid. The DNA sequence was analyzed by DNA sequencing (Cogenics). Next, the pRD4 was transformed in *E.coli* DH5α to obtain *E.coli* DH5α (*luxS*) strain.

### Labeling of *D. vulgaris* with calcein-acetoxymethyl-ester (AM)

The labelling of *D. vulgaris* cells was carried out as described by Benomar *et al*.^18^. Briefly, *D. vulgaris* cells were grown in Starkey medium^56^ under anaerobic conditions, then exponentially growing *D. vulgaris* cells (5 ml) was harvested at room temperature by centrifugation at 4000g for 10 min, washed twice with Starkey medium and resuspended in 5 ml fresh Starkey medium. 100 μl of calcein-AM (1 mg/ml in dimethylsulfoxide) were then added to the medium. The suspension was incubated in the dark at 37°C for 2 hours under anaerobic conditions. Cells were subsequently harvested and washed three times in fresh, dye-free GY medium and used in the exchange experiments.

### Labelling of *E. coli* DH5α and *E. coli* DH10 with mCherry

The labelling was carried out as described by Benomar *et al*^18^.

### Exchange of cytoplasmic molecules between *D. vulgaris* and *E. coli*

To study the exchange of molecules between the two bacteria, *D. vulgaris* labelled with calcein was mixed with several strains of *E. coli* labelled with mCherry: *E. coli* DH10B strain, producer of AI-2 molecule, *E. coli* DH5α non producer of AI-2 molecule, DH5α (*lux*S). In some cases, unlabelled cells of *D. vulgaris* were mixed with *E. coli* labelled with mCherry. The mixture was diluted in 5 ml fresh GY medium and put in a tube containing a coverslip and incubated at 37°C for 20 h. The coverslip was removed after 20 h of growth and bacterial cells attached to coverslip were visualized by fluorescence confocal microscopy as previously described in Benomar *et al*^18^.

### AI-2 activity assay

AI-2 activities of cell-free culture supernatants were measured by using *Vibrio harveyi* reporter strain BB170 as described by Bassler *et al*^59^. Briefly, an overnight culture of *V. harveyi* (grown for 16 h in AB medium) was diluted 1/5,000 in fresh AB medium (300 mM NaCl, 50 mM MgSO_4_, 2% [wt/v] Casamino Acids, 10 mM potassium phosphate [pH 7], 1 mM L-arginine, 1% [wt/v] glycerol). The diluted cells (90 μL) were added to 96-well plates (Corning) containing 10 μL of the cell-free culture supernatant of *E. coli* or *D. vulgaris* or *C. acetobutylicum*, obtained after centrifugation and filtration through 0.2m membranes to remove bacterial cells, or synthetic AI-2 molecule. The microtiter plate was incubated at 30 °C with shaking at 160 rpm and bioluminescence was measured each hour over the course of 6 h using a Tecan GENioS plate reader (Tecan, USA). AI-2 activity is reported as fold induction of bioluminescence over background. The reported values represent the average bioluminescence stimulated by three independent preparations of cell-free culture fluids or synthetic AI-2 molecule. Similar experiments were performed in presence of various amounts of *D. vulgaris* culture supernatant in Starkey medium.

### Use of RedoxSensor Green as a probe for active respiration in *D. vulgaris*

RedoxSensor Green (RSG) (Backlight™ RedoxSensor™ Green Vitality Kit, Life Technologies) was used to assess cellular respiration activity of *D. vulgaris. D. vulgaris* cells were taken from Starkey medium culture in mid-log phase, washed twice and starved by incubation in GY medium (without lactate and sulfate) for 20 hours at 37°C. Following starvation, *D. vulgaris* cells were harvested at room temperature by centrifugation at 4,000 g for 10 min, washed twice with GY medium and diluted in 5 ml fresh GY medium containing 1 μM RSG reagent. The cultures were supplemented with 10 mM lactate and 10 mM sulfate or not and bacterial cells were imaged by fluorescence microscopy following incubation at 37°C for 20h. After 20h of incubation at 37 °C, the co-culture was left for 5 min in contact with FM4-64 to visualize the bacterial membrane and to be able to see *C. acetobutylicum*, which was not labelled at the beginning of the experiment. Also, the effect of the electron transport chain uncoupler CCCP on RSG fluorescence was verified to further confirm the redox sensing functionality of RSG in *D. vulgaris* as previously reported for other bacteria^9^.

### Analysis and purification of the Antagonist AI-2 compounds from *D. vulgaris*

*D. vulgaris* cells were grown in Starkey medium under anaerobic conditions for 30h at 37 °C. The culture was then centrifuged at 10,000 rpm for 5 min and filtered through 0.2 μm membranes to remove the cells. The cell-free supernatants were stored at −20°C or immediately analysed by HPLC. The analysis was carried out on an Agilent 1200 HPLC system equipped with a UV detector and a refractometer (Agilent technologies). Separation (20-50μl of samples are injected) was achieved on an Agilent POROSHELL EC-C18 reverse-phase column (C18, 4.6×150 mm, 2.7μm) set at 30°C. The compounds were eluted with 8 % solution A (Acetonitrile + 1 %(v/v) formic acid) and 92 % of solution B (Deionized water + 1%(v/v) formic acid), at a flow rate of 0.6 ml/min.

### Carbon exchange by Stable isotope probing (SIP)

SIP method described by Neufeld *et al^60^* and derived from Meselson and Stahl was slightly modified. Glucose-yeast extract (GY) medium was prepared with D-glucose-^13^C6 (14 mM) from Cortecnet and 0.1% yeast extract (^12^C), supplemented with similar inorganic nutrients as used for the Starkey preparation (MgCl_2_ instead of MgSO_4_) and N_2_ in the headspace.

^13^C-GY medium was inoculated (10%) with either washed *C. acetobutylicum* enriched in ^13^C *via* 26 subculturing or *D. vulgaris* ^12^C at exponential growth phase, or with the two strains, to constitute an artificial consortium in a 1:1 ratio according to the absorbance at 600 nm.

At the end of the exponential phase, genomic DNA was extracted from the cell pellets using the NucleoBond AXG20 kit (Macherey-Nagel), and DNA purity and concentration were determined with the Nanodrop 2000c spectrophotometer (Thermoscientific).

To separate labelled/heavier (^13^C-DNA) from unlabelled/lighter (^12^C-DNA), density gradient centrifugation was performed in 5,1 ml quick-seal tubes in a NVT 65.2 rotor (near vertical) using Optima L-90K centrifuge (Beckman coulter). CsCl medium having an average density of 1,72g/ml was loaded with 6μg of total extracted DNA. After centrifugation at 20°C for 66h at 41500 rpm 169000 g_av_, each gradient (^13^C-labelled-*C. acetobutylicum* + ^12^C-unlabelled-*D. vulgaris* and ^12^C-unlabelled-*C. acetobutylicum* + ^12^C-unlabelled-*D. vulgaris) was* fractioned from the bottom to the top by displacement with mineral oil. DNA in each fraction was precipitated as described by Neufeld *et al*^61^.

The presence and relative amount of DNA from each bacterium were followed. The primers used for the qPCR (*dsrA* and *endoG*) are listed in Supplementary Table 1. The reaction was performed with GoTaq mix and the PCR was carried out in a Techne Prime Elite thermal cycler as follows: 2min at 98°C for the initial activation of enzymes, 21 cycles of 30s at 98°C, 30s at 58°C and 2 min at 72°C. Experiments were made in triplicate.

### AI-2 synthesis

AI-2 was obtained in a four step sequence starting from the commercially available methyl (S)-(-)-2,2-dimethyl-1,3-dioxolane-4-carboxylate adapting the reported procedures^62,63^.

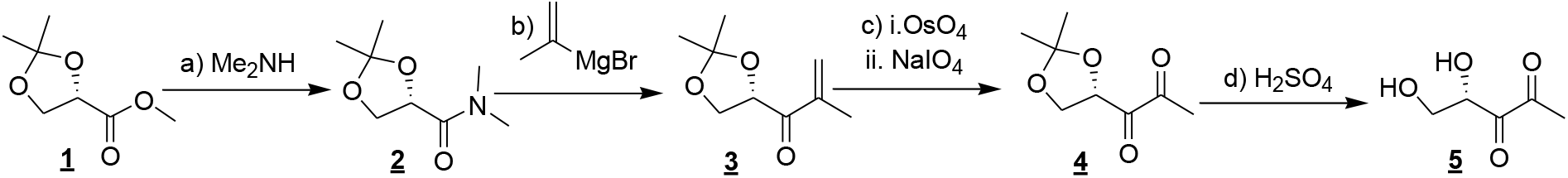

**<u>Scheme</u>** : a) Me_2_NH, EtOH, 0°C - RT, 48hrs, 86%; b) *i*-PropenylMgBr, Et_2_O 48hrs, 46%; c) i.OsO_4_, NMO, CH_3_COCH_3_ : H_2_O; ii.NaIO_4_, MeOH : H_2_O, RT, 30 min, 25% over 2 steps; d) H_2_SO_4_, D_2_O : d_6_-DMSO, 0 °C, 1hrs.

First, the methyl ester (<u>**1)**</u> was transformed into the amide (<u>**2)**</u> by reacting with dimethylamine in EtOH. The reaction of amide (<u>**2)**</u> with isopropenyl magnesium bromide gives olefin (<u>**3)**</u>. Dihydroxylation of olefin (<u>**3)**</u> with catalytic osmium tetroxide in the presence of N-methylmorpholine-N-oxide and subsequent cleavage of generated diol with NaIO_4_ produced ketone (<u>**4)**</u>. Finally, the hydrolysis of dioxolane ring in acidic condition yields AI-2 (<u>**5**</u>) and its cyclic anomeric products. Spectral data were consistent with those previously reported. Details of each step are detailed below.

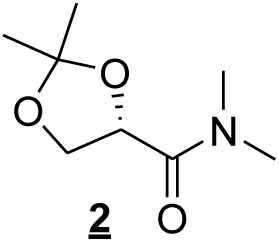

#### *N,N*-Dimethyl (*S*)-α,β-isopropylidene glyceramide (<u>2</u>)

To the solution containing dimethylamine (20 mL, 30 vol. % in ethanol) was added methyl (*S*) - α, β-isopropylidene glycerate (3g, 18.5 mmol) at 0 °C. The mixture was stirred for 24 h. After adding an additional 10 mL of the dimethylamine solution, the stirring was continued for further 24 h. The volatile compounds were evaporated and the residue was purified by column chromatography to yield 2.8g of amide <u>**2**</u> (86%, colorless liquid); R*_f_* = 0.2 (pentane / ethyl acetate, 3:2); ^1^H NMR (400 MHz, CDCl_3_) ∂ = 4.62 (t, *J* = 6.65 Hz, 1H), 4.31 (dd, *J* = 8.28, 6.53 Hz, 1H), 4.07 (dd, *J* = 8.53, 6.78 Hz, 1H), 3.05 (s, 3H), 2.90 (s, 3H), 1.34 (s, 6H).

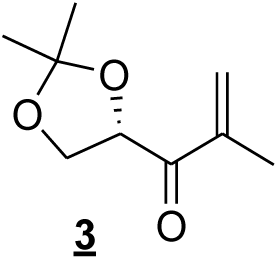

#### (*S*)-4-Methacryloyl-2,2-dimethyl-1,3-dioxolane (<u>3</u>)

To a solution of amide <u>**2**</u> (1.74g, 10 mmol) in anhydrous diethyl ether(10 mL) at 0 °C under an Argon atmosphere, 21.0 mL of a 0.5M solution of isopropenylmagnesium bromide in THF was added and stirred for 15 minutes. The reaction mixture was quenched with 1M HCl and extracted 3 times with diethyl ether and the combined organic phase dried over Na_2_SO_4_ and evaporated to dryness. The residue was purified by chromatography on SiO_2_ to afford <u>**3**</u> (0.8 g, 46 %). R*_f_* = 0.33 (pentane / ethyl acetate, 9:1); ^1^H NMR (300 MHz, CDCl_3_) *δ=* 6.06 (s, 1 H), 5.91 (d, *J* = 1.38, 1 H), 5.08 (dd, *J* = 7.26,6.03, 1 H), 4.22 (dd, *J* = 8.34, 7.4, 1 H), 4.09 (dd, *J* = 8.43,5.94, 1 H), 1.89 (s, 3 H), 1.41 (s, 6 H).

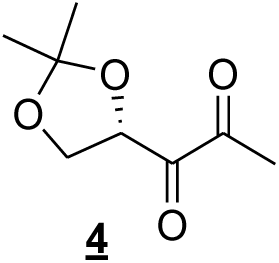

#### 1-(2,2-Dimethyl-[1,3]dioxolan-4-yl)-propane-1,2-dione (<u>4</u>)

The alkene (<u>**3**</u>) (40 mg, 0.23 mmol) was dissolved in a mixture of acetone/water (4mL/1mL). *N*-Methyl morpholine-*N*-oxide monohydrate (39 mg, 0.23 mmol) was added slowly and the reaction mixture was stirred at room temperature for 10 min. Then 4% aqueous solution of osmium tetraoxide (0.1 mL, 0.011 mmol) was added to the above reaction mixture and stirred overnight at room temperature. The mixture was quenched with Na_2_SO_3_ and extracted with dichloromethane. The organic layer was dried (Na_2_SO_4_), filtered and the filtrate concentrated under reduced pressure. The crude diol was used in next step as such. To the diol in methanol (3.5mL) and water (1.5mL) was added sodium periodate (0.146g, 0.69 mmol) and stirred at RT for 30min. The reaction mixture was diluted with water and extracted with dichloromethane. The organic layer was dried (Na_2_SO_4_), filtered and the filtrate was concentrated under reduced pressure. The residue was purified by chromatography on SiO_2_ to afford <u>**4**</u> (15.9 mg, 40 % over 2 steps). R*_f_* = 0.65 (pentane / ethyl acetate, 6:4); ^1^H NMR (400 MHz, CDCl_3_) *δ*= 5.14 (dd, *J* = 7.8, 5.8, 1 H), 4.37 (t, *J* = 8.4, 1 H), 4.0 (dd, *J* = 9.00, 5.4, 1 H), 2.39 (s, 3 H), 1.47 (s, 3 H), 1.42 (s, 3 H).

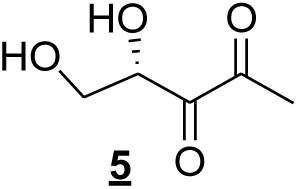

#### (*S*)-4, 5-Dihydroxy-pentane-2, 3-Dione (<u>5</u>)

To a solution of diketone <u>**4**</u> (3.0 mg, 0.017mmol) in D2O (0.5 mL) and *d*_6_-DMSO (0.2 mL) under an Argon atmosphere, H_2_SO_4_ (0.005 mL) was added and stirred at RT for 1 h. The reaction mixture was quenched with 5 mg Na_2_CO_3_, filtered to get 0.0249 M solution of AI-2 (<u>**5**</u>) and its cyclic anomeric products in D_2_O and *d*_6_-DMSO. Spectral data were consistent with those previously reported. ^1^H-NMR (300 MHz, D2O): 4.25 (t, *J*= 6.4 Hz, 1H), 4.10-4.02 (m, 2H), 3.93 (dd, *J*= 3.4, 5.6 Hz,1H), 3.76-3.67 (m, 2H), 3.84-3.81 (m, 1H), 3.53 (dd, *J*= 7.5, 11.6 Hz, 1H), 3.45 (dd, *J*=5.4, 9.2 Hz, 1H), 2.27 (s, 3H), 1.34 (s, 3H), 1.31 (s, 3H). HRMS (ESI-TOF) m/z: [M - H] ^-^ calcd for C5H7O_4_^-^ 131.0350; found 131.0354.

## Acknowledgements

We acknowledge Pascale Infossi and Dr Deborah Byrne for help with some experiments. This work has benefited from the facilities and expertise of the Platform for Microscopy of IMM. We thank Dr Jean-Philippe Steyer and Dr Chantal Iobbi for helpful discussions. Dr A. Cornish-Bowden for helpful discussions and correction of the English. This work was supported by the A*MIDEX project (nANR-11-IDEX-0001-02) from Aix Marseille University.

## Author contributions

DR, C.B, OO and MTGO designed the research; DR, CB, and GK performed the research; DR, CB, MG, AS and MTGO. analyzed the data; and DR, MLC. and MTGO wrote the paper. DR and CB contributed equally to this work. The authors declare no competing interests.

## Supplementary information and figures

**Supplementary Table 1:**
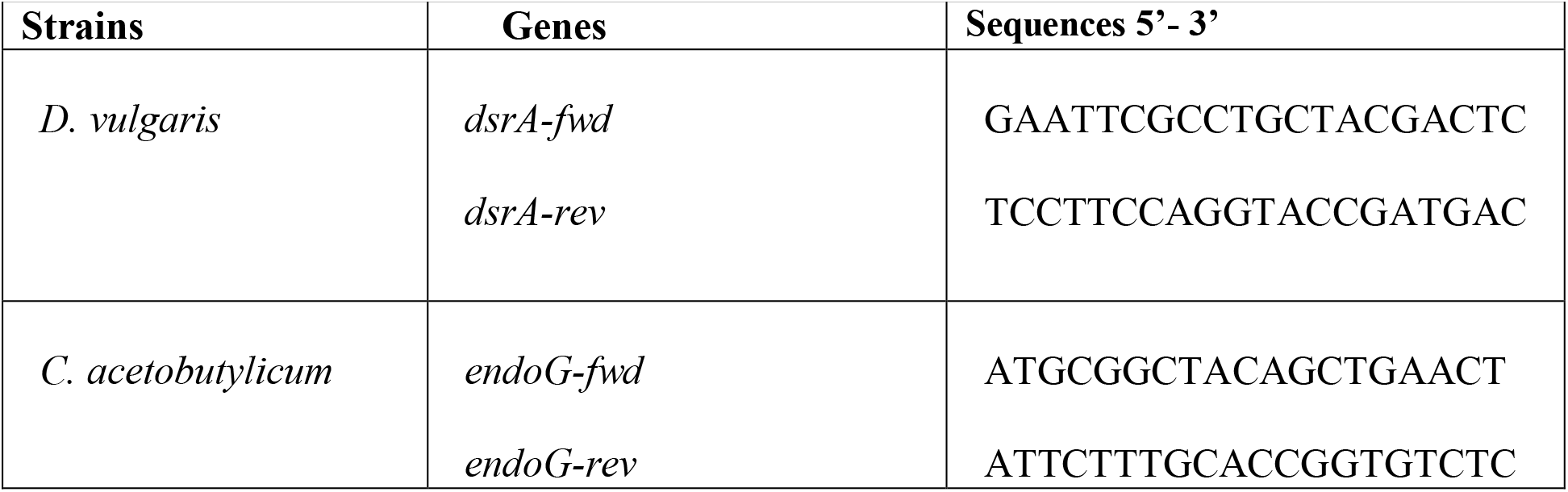
Primers used for PCR quantification for *D. vulgaris* and *C. acetobutylicum*.

## Supplementary Figure Legends

**Supplementary Figure 1:**
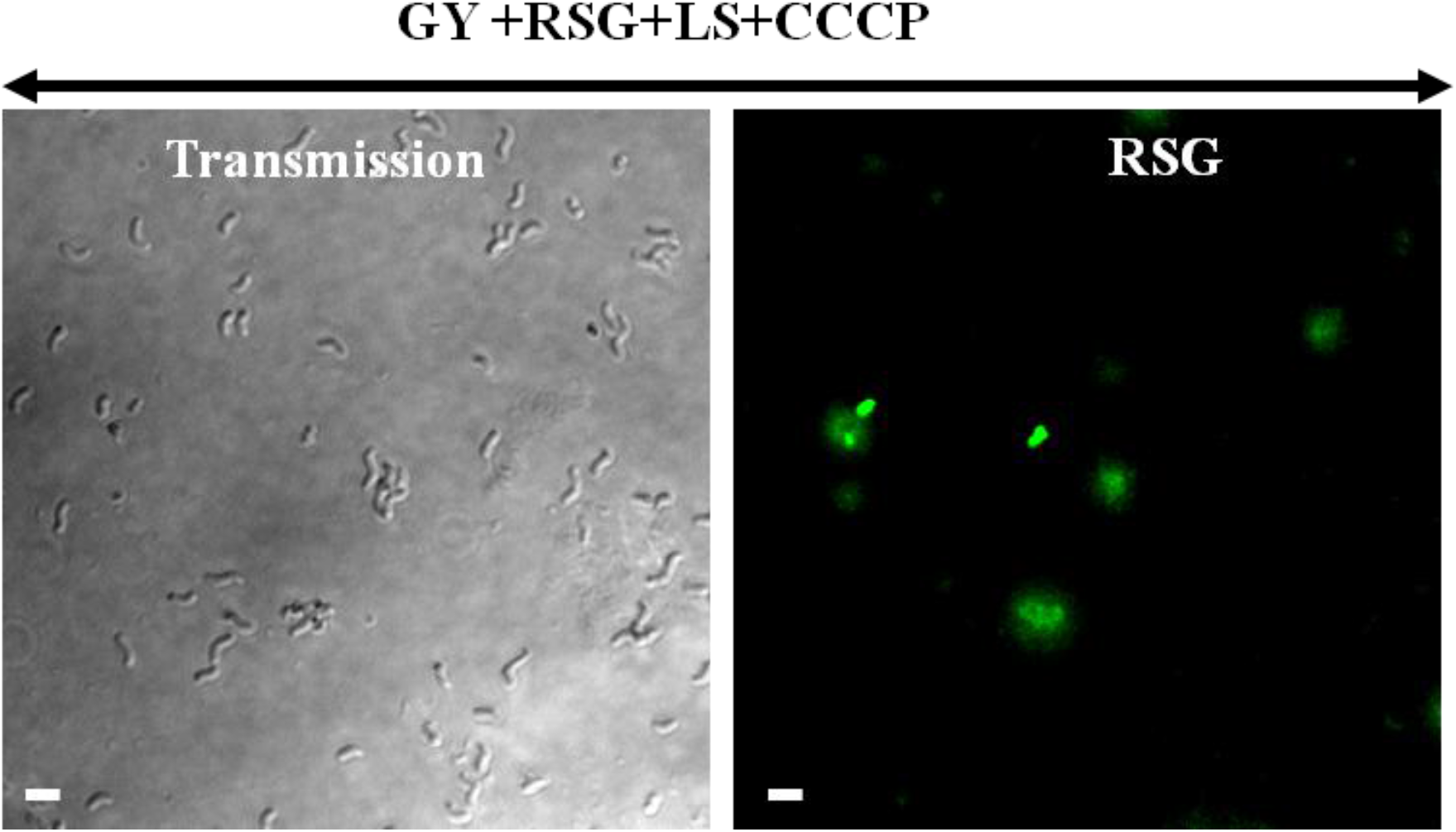
The effect of electron transport chain uncoupler on RedoxSensor Green fluorescence. *D. vulgaris* growing exponentially in Starkey medium was washed twice and starved by incubation in GY medium, 20h at 37°C. A starved culture was diluted in GY medium 5 ml containing 10mM of lactate/sulfate and supplemented with 1μM RSG and 10μM uncoupler CCCP (final concentrations). The culture was sampled and visualized by fluorescence confocal microscopy after 20h incubation at 37 °C. Scale bar, 2μm in all panels. (n=3).

**Supplementary Figure 2:**
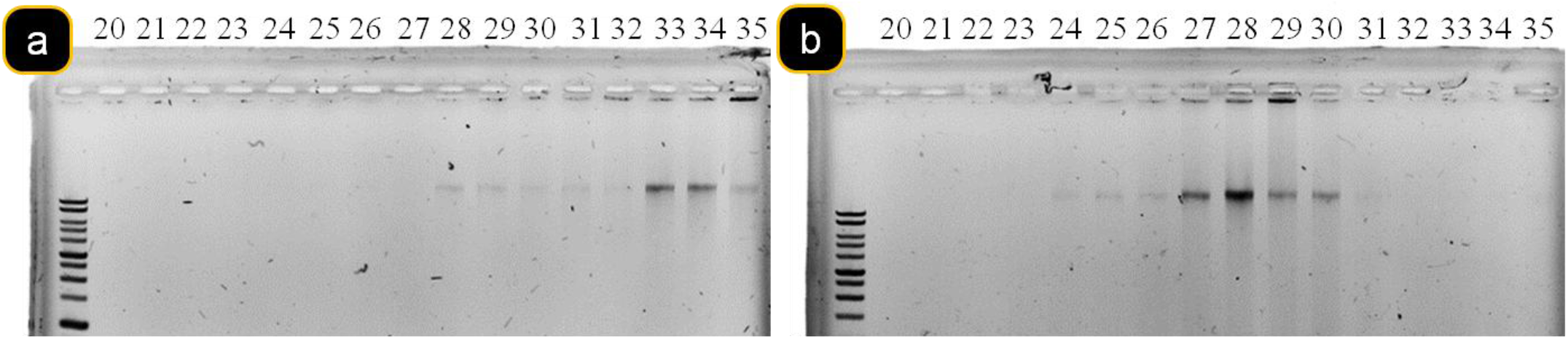
Analysis of fractions from density gradient on agarose gel. Fractions of *C. acetobutylicum* ^12^C + *D. vulgaris* ^12^C in ^12^C glucose medium DNA as negative control (a) and *C. acetobutylicum* ^13^C + *D. vulgaris* ^12^C in ^13^C glucose medium DNA (b). a and b were centrifuged together. Fractions 27 to 30 contained ^13^C DNA (heavier) and fraction 33 to 35 contained ^12^C DNA (lighter).

**Supplementary Figure 3:**
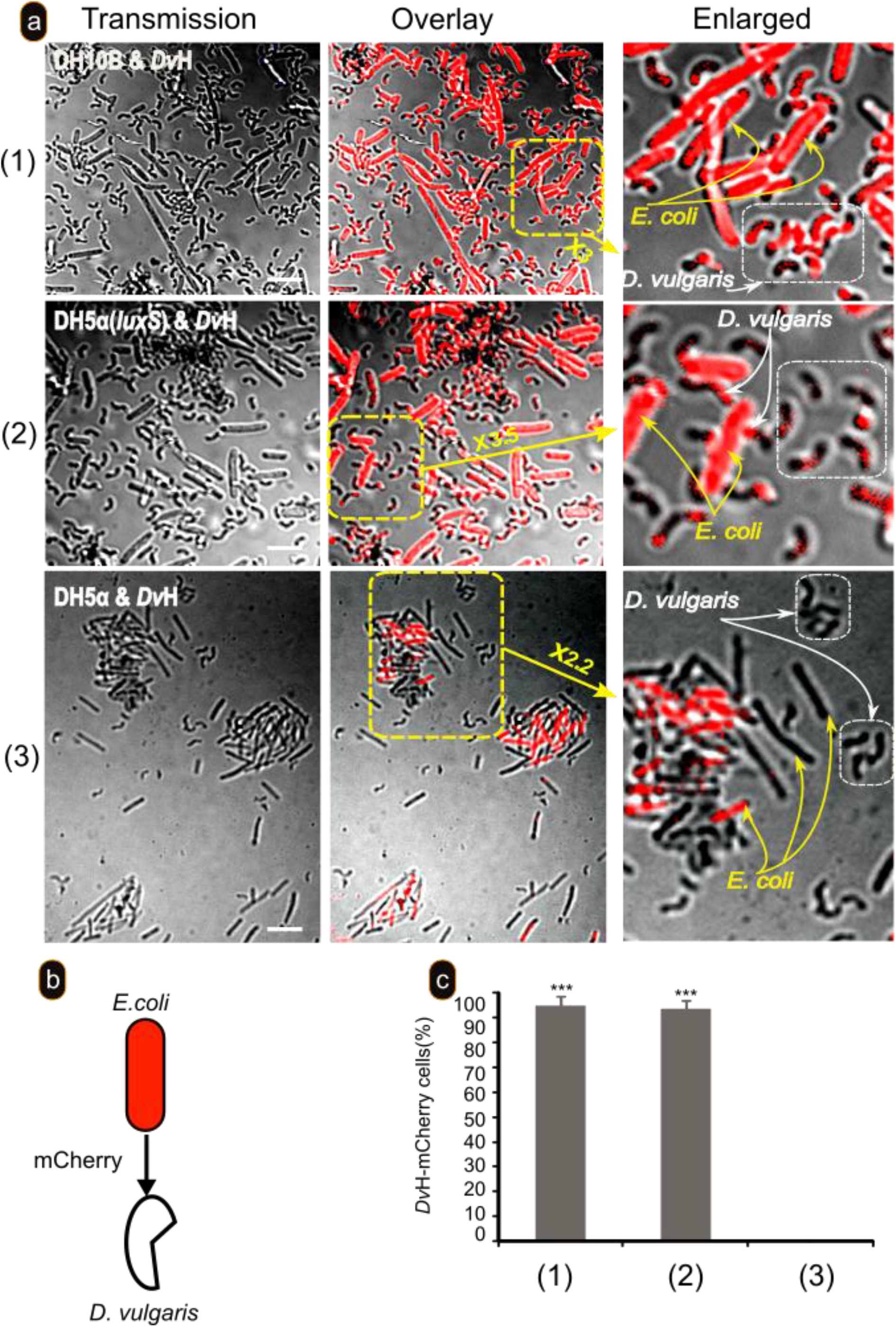
AI-2 is required for mCherry transfer from *E. coli* to *D. vulgaris*. *D. vulgaris* (*Dv*H) growing exponentially in Starkey medium was washed in GY medium, mixed *E. coli* strains DH10B, DH5α and DH5α (*luxS*) labelled with mCherry and visualized by fluorescence confocal microscopy after 20-h incubation at 37°C in GY medium. Scale bar, 2μm in all panels.

**Supplementary Figure 4:**
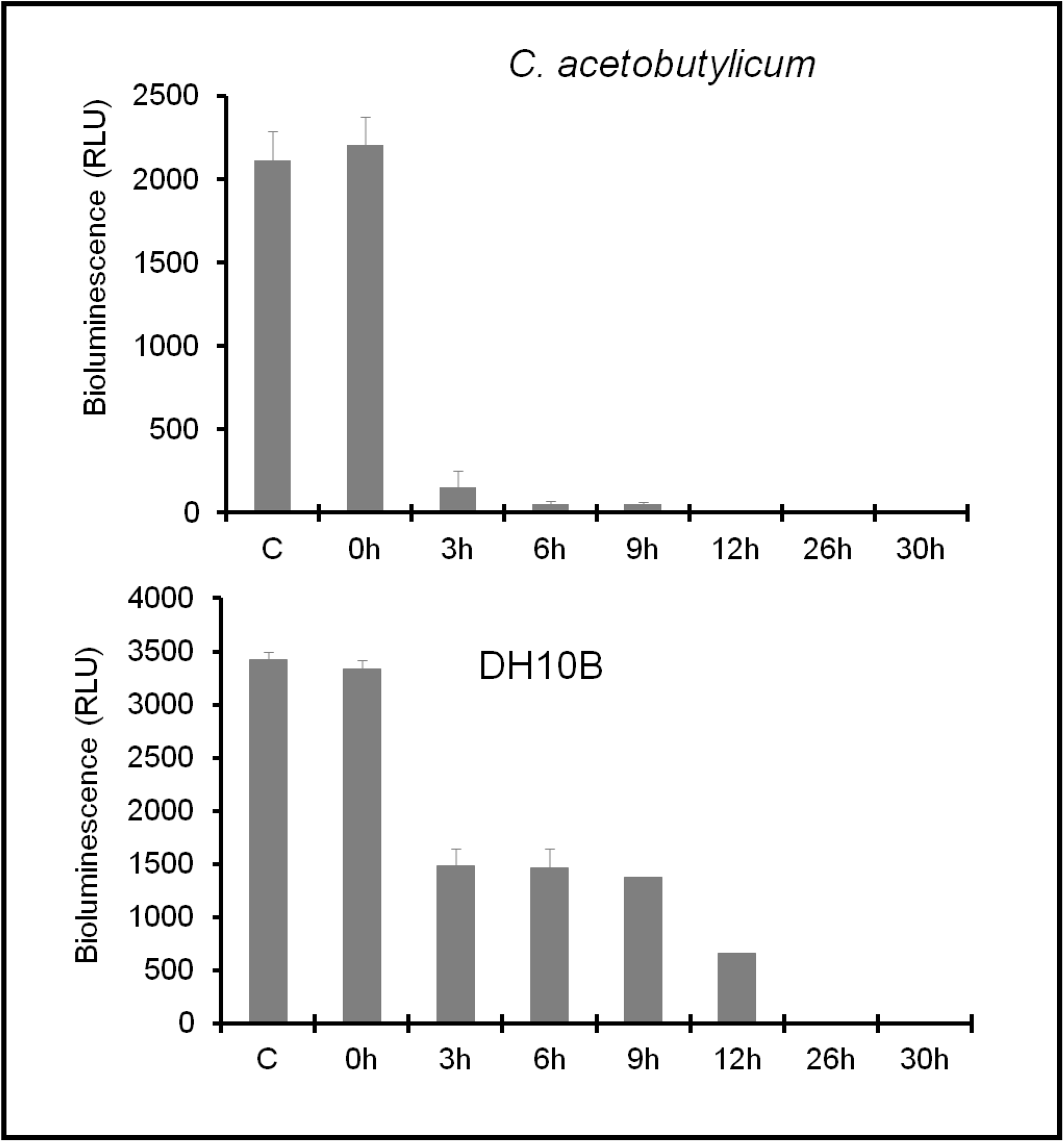
Kinetics of production of the inhibitors of AI-2 activity by *D. vulgaris* in Starkey medium. Supernatants from pure cultures of *C. acetobutylicum* and *E. coli* DH10B were taken after 30 hours of culture in GY medium and filtered (0.2μm). Then, the activity of AI-2 in the filtered samples was analyzed using *V. harveyi* reporter strain BB170 in the absence (C) or in the presence of 5μL of *D. vulgaris* supernatant (grown in Starkey medium) taken at different growth times. (n=3).

**Supplementary Figure 5:**
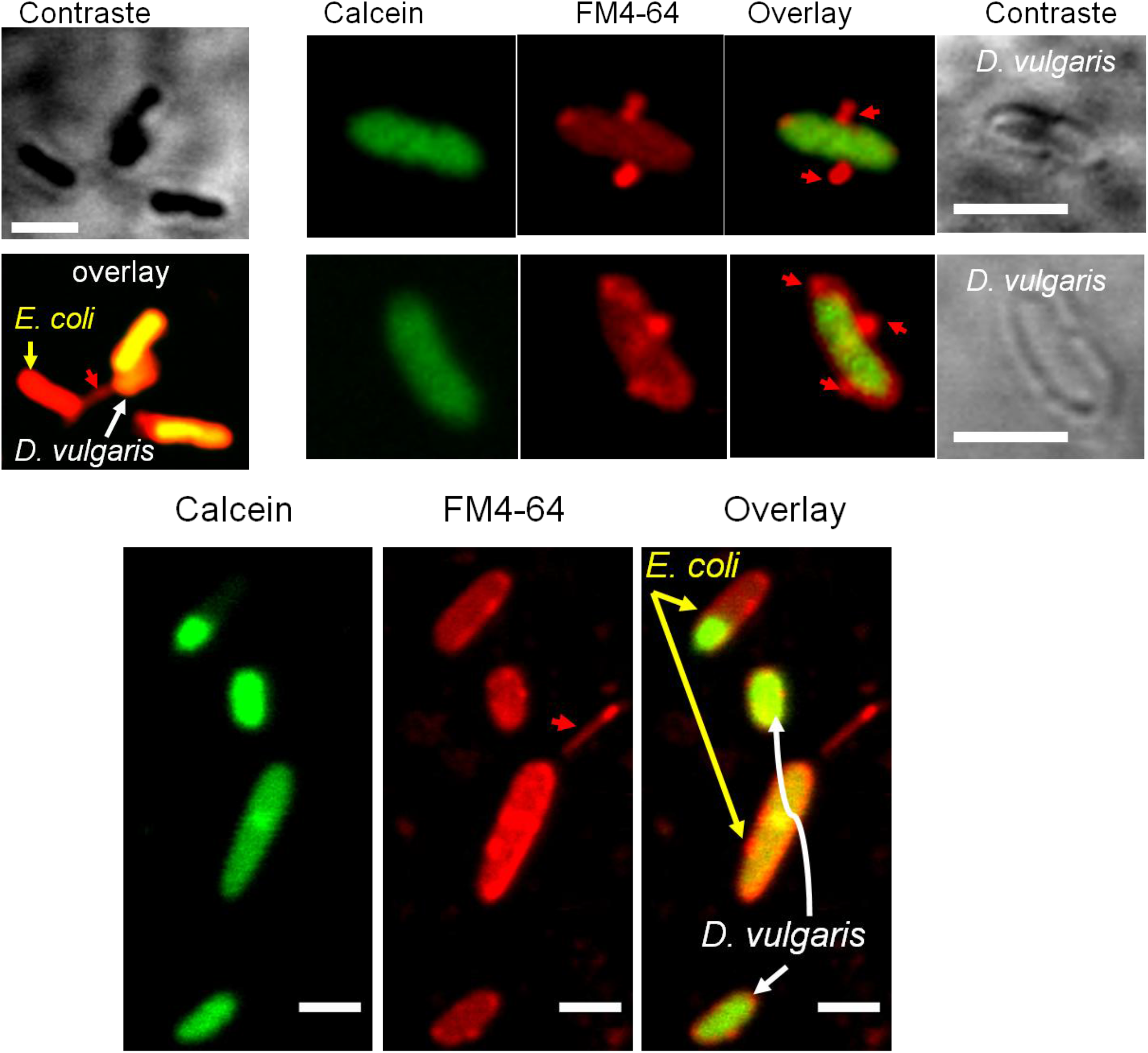
*D. vulgaris* growing exponentially in Starkey medium was labelled with calcein, washed with GY medium, and mixed with unlabelled *E. coli* strains DH10B. After 20h incubation at 37 °C, the culture was sampled, left for 5 min in contact with FM4-64 (in order to visualize the two strains) and visualized by fluorescence confocal microscopy Red arrows indicate apparent connections between bacterial cells. Scale bar, 2 μm in all panels.

## Notes

### Competing Interest Statement

The authors have declared no competing interest.

## Bibliography

1 Conrad, R. Soil microorganisms as controllers of atmospheric trace gases (H2, CO, CH4, OCS, N2O, and NO). Microbiol Rev 60, 609–640 (1996).

2 Newman, D. K. & Banfield, J. F. Geomicrobiology: how molecular-scale interactions underpin biogeochemical systems. Science 296, 1071–1077, doi:10.1126/science.1010716 (2002).

3 Falkowski, P. G., Fenchel, T. & Delong, E. F. The microbial engines that drive Earth’s biogeochemical cycles. Science 320, 1034–1039, doi:10.1126/science.1153213 (2008).

4 Gibbons, S. M., Gilbert, J.A. Microbial diversity--exploration of natural ecosystems and microbiomes. Curr Opin Genet Dev. 35, 66–72, doi:10.1016/j.gde.2015.10.003 (2015).

5 Haruta, S., Kato, S., Yamamoto, K. & Igarashi, Y. Intertwined interspecies relationships: approaches to untangle the microbial network. Environ Microbiol 11, 2963–2969, doi:10.1111/j.1462-2920.2009.01956.x (2009).

6 Phelan, V. V., Liu WT, Pogliano K, Dorrestein PC. Microbial metabolic exchange—the chemotype-to-phenotype link. Nat Chem Biol 8, 26–35, doi:10.1038/nchembio.739 (2011).

7 Little, A. E., Robinson, C. J., Peterson, S. B., Raffa, K. F. & Handelsman, J. Rules of engagement: interspecies interactions that regulate microbial communities. Annu Rev Microbiol 62, 375–401, doi:10.1146/annurev.micro.030608.101423 (2008).

8 Morris, J. J. Black Queen evolution: the role of leakiness in structuring microbial communities. Trends Genet 31, 475–482, doi:10.1016/j.tig.2015.05.004 (2015).

9 Pirbadian, S. et al. Shewanella oneidensis MR-1 nanowires are outer membrane and periplasmic extensions of the extracellular electron transport components. Proc Natl Acad Sci USA 111, 12883–12888, doi:10.1073/pnas.1410551111 (2014).

10 Dubey, G. P. & Ben-Yehuda, S. Intercellular nanotubes mediate bacterial communication. Cell 144, 590–600, doi:10.1016/j.cell.2011.01.015 (2011).

11 Dubey, G. P. et al. Architecture and Characteristics of Bacterial Nanotubes. Dev Cell 36, 453–461, doi:10.1016/j.devcel.2016.01.013 (2016).

12 Mamou, G., Malli Mohan, G. B., Rouvinski, A., Rosenberg, A. & Ben-Yehuda, S. Early Developmental Program Shapes Colony Morphology in Bacteria. Cell Rep 14, 1850–1857, doi:10.1016/j.celrep.2016.01.071 (2016).

13 D’Souza, G. et al. Ecology and evolution of metabolic cross-feeding interactions in bacteria. Nat Prod Rep 35, 455–488, doi:10.1039/c8np00009c (2018).

14 D’Souza, G. & Kost, C. Experimental Evolution of Metabolic Dependency in Bacteria. PLoS Genet 12, e1006364, doi:10.1371/journal.pgen.1006364 (2016).

15 De Roy, K., Marzorati, M., Van den Abbeele, P., Van de Wiele, T. & Boon, N. Synthetic microbial ecosystems: an exciting tool to understand and apply microbial communities. Environ Microbiol 16, 1472–1481, doi:10.1111/1462-2920.12343 (2014).

16 Heidelberg, J. F. et al. The genome sequence of the anaerobic, sulfate-reducing bacterium Desulfovibrio vulgaris Hildenborough. Nat Biotechnol 22, 554–559, doi:10.1038/nbt959 (2004).

17 Muyzer, G. & Stams, A. J. The ecology and biotechnology of sulphate-reducing bacteria. Nat Rev Microbiol 6, 441–454, doi:10.1038/nrmicro1892 (2008).

18 Benomar, S. et al. Nutritional stress induces exchange of cell material and energetic coupling between bacterial species. Nature Communications 6, doi:10.1038/ncomms7283 (2015).

19 Walker, C. B. et al. Functional responses of methanogenic archaea to syntrophic growth. Isme Journal 6, 2045–2055, doi:10.1038/ismej.2012.60 (2012).

20 Pande, S. et al. Metabolic cross-feeding via intercellular nanotubes among bacteria. Nature Communications 6, doi:10.1038/ncomms7238 (2015).

21 Pande, S. & Kost, C. Bacterial Unculturability and the Formation of Intercellular Metabolic Networks. Trends Microbiol 25, 349–361, doi:10.1016/j.tim.2017.02.015 (2017).

22 Pande, S. & Velicer, G. J. Chimeric Synergy in Natural Social Groups of a Cooperative Microbe. Curr Biol 28, 487, doi:10.1016/j.cub.2018.01.044 (2018).

23 Baidya, A. K., Bhattacharya, S., Dubey, G. P., Mamou, G. & Ben-Yehuda, S. Bacterial nanotubes: a conduit for intercellular molecular trade. Curr Opin Microbiol 42, 1–6, doi:10.1016/j.mib.2017.08.006 (2018).

24 Abisado, R. G., Benomar, S., Klaus, J. R., Dandekar, A. A. & Chandler, J. R. Bacterial Quorum Sensing and Microbial Community Interactions. MBio 9, doi:10.1128/mBio.02331-17 (2018).

25 Kalyuzhnaya, M. G., Lidstrom, M. E. & Chistoserdova, L. Real-time detection of actively metabolizing microbes by redox sensing as applied to methylotroph populations in Lake Washington. Isme Journal 2, 696–706, doi:10.1038/ismej.2008.32 (2008).

26 Bassler, B. L. & Losick, R. Bacterially speaking. Cell 125, 237–246, doi:10.1016/j.cell.2006.04.001 (2006).

27 Vendeville, A., Winzer, K., Heurlier, K., Tang, C. M. & Hardie, K. R. Making ‘sense’ of metabolism: autoinducer-2, LuxS and pathogenic bacteria. Nat Rev Microbiol 3, 383–396, doi:10.1038/nrmicro1146 (2005).

28 Waters, C. M. & Bassler, B. L. Quorum sensing: Cell-to-cell communication in bacteria. Annual Review of Cell and Developmental Biology 21, 319–346, doi:10.1146/annurev.cellbio.21.012704.131001 (2005).

29 Kawaguchi, T., Chen, Y. P., Norman, R. S. & Decho, A. W. Rapid screening of quorum-sensing signal N-acyl homoserine lactones by an in vitro cell-free assay. Appl Environ Microbiol 74, 3667–3671, doi:10.1128/AEM.02869-07 (2008).

30 Chen, X., Schauder, S., Potier, N., Van Dorsselaer, A., Pelczer, I., Bassler, B.L. and Hughson, F.M. Structural identification of a bacterial quorum-sensing signal containing boron. Nature 415, 545–549 (2002).

31 Roy, V., Smith; JA, Wang; J, Stewart: JE, Bentley: WE, Sintim: HO. Synthetic Analogs Tailor Native AI-2 Signaling Across Bacterial Species. J. Am. Chem. Soc. 132 11141–11150, doi:10.1021/ja102587w (2010).

32 Wang, Y., Cui, J., Sun, X., Zhang, Y. Tunneling-nanotube development in astrocytes depends on p53 activation. Cell Death Differ. 18, 732–742, doi: 10.1038/cdd.2010.147 (2011).

33 Stephens, K., Pozo, M., Tsao, C-Y., Hauk, P., Bentley, W, E. Bacterial co-culture with cell signaling translator and growth controller modules for autonomously regulated culture composition. Nature Communications 10, doi:10.1038/s41467-019-12027-6 (2019).

34 Yong, Y. C., Wu, X. Y., Sun, J. Z., Cao, Y. X. & Song, H. Engineering quorum sensing signaling of Pseudomonas for enhanced wastewater treatment and electricity harvest: A review. Chemosphere 140, 18–25, doi:10.1016/j.chemosphere.2014.10.020 (2015).

35 Shrout, J. D. & Nerenberg, R. Monitoring bacterial twitter: does quorum sensing determine the behavior of water and wastewater treatment biofilms? Environ Sci Technol 46, 1995–2005, doi:10.1021/es203933h (2012).

36 Pereira, C. S., McAuley, J. R., Taga, M. E., Xavier, K. B. & Miller, S. T. Sinorhizobium meliloti, a bacterium lacking the autoinducer-2 (AI-2) synthase, responds to AI-2 supplied by other bacteria. Molecular Microbiology 70, 1223–1235, doi:10.1111/j.1365-2958.2008.06477.x (2008).

37 Brito, P. H., Rocha, E. P., Xavier, K. B. & Gordo, I. Natural genome diversity of AI-2 quorum sensing in Escherichia coli: conserved signal production but labile signal reception. Genome Biol Evol 5, 16–30, doi:10.1093/gbe/evs122 (2013).

38 Duan, K. M., Dammel, C., Stein, J., Rabin, H. & Surette, M. G. Modulation of Pseudomonas aeruginosa gene expression by host microflora through interspecies communication. Molecular Microbiology 50, 1477–1491, doi:10.1046/j.1365-2958.2003.03803.x (2003).

39 Pereira, C. S., de Regt, A. K., Brito, P. H., Miller, S. T. & Xavier, K. B. Identification of Functional LsrB-Like Autoinducer-2 Receptors. Journal of Bacteriology 191, 6975–6987, doi:10.1128/jb.00976-09 (2009).

40 Anderson, J. K. et al. Chemorepulsion from the Quorum Signal Autoinducer-2 Promotes Helicobacter pylori Biofilm Dispersal. MBio 6, e00379, doi:10.1128/mBio.00379-15 (2015).

41 Hense, B. A. & Schuster, M. Core principles of bacterial autoinducer systems. Microbiol Mol Biol Rev 79, 153–169, doi:10.1128/MMBR.00024-14 (2015).

42 Bari, S. M., Roky, M.K., Mohiuddin, M., Kamruzzaman, M., Mekalanos, J.J, Faruque, S.M. Quorum-sensing autoinducers resuscitate dormant Vibrio cholerae in environmental water samples. Proc Natl Acad Sci U S A 110 9926–9931, doi:10.1073/pnas.1307697110 (2013).

43 Ayrapetyan, M., Williams, T.C., Oliver, J.D. Interspecific quorum sensing mediates the resuscitation of viable but nonculturable vibrios. Appl Environ Microbiol. 80, 2478–2483, doi:10.1128/AEM.00080-14 (2014).

44 Boyle, K. E., Monaco, H., van Ditmarsch, D., Deforet, M. & Xavier, J. B. Integration of Metabolic and Quorum Sensing Signals Governing the Decision to Cooperate in a Bacterial Social Trait. PLoS Comput Biol 11, e1004279, doi:10.1371/journal.pcbi.1004279 (2015).

45 Reimmann, C. et al. Genetically programmed autoinducer destruction reduces virulence gene expression and swarming motility in Pseudomonas aeruginosa PAO1. Microbiology-Sgm 148, 923–932 (2002).

46 Zhang, H. B., Wang, L. H. & Zhang, L. H. Genetic control of quorum-sensing signal turnover in Agrobacterium tumefaciens. Proc Natl Acad Sci U S A 99, 4638–4643, doi:10.1073/pnas.022056699 (2002).

47 Rasmussen, T. B. et al. Identity and effects of quorum-sensing inhibitors produced by Penicillium species. Microbiology-Sgm 151, 1325–1340, doi:10.1099/mic.0.27715-0 (2005).

48 Chu, Y. Y. et al. A new class of quorum quenching molecules from Staphylococcus species affects communication and growth of gram-negative bacteria. PLoS Pathog 9, e1003654, doi:10.1371/journal.ppat.1003654 (2013).

49 Singh, V. et al. Structure and inhibition of a quorum sensing target from Streptococcus pneumoniae. Biochemistry 45, 12929–12941, doi:10.1021/bi061184i (2006).

50 Shen, G., Rajan, R., Zhu, J., Bell, C.E., Pei, D. Design and synthesis of substrate and intermediate analogue inhibitors of S-ribosylhomocysteinase. J Med Chem. 49, 3003–3011, doi:10.1021/jm060047g (2006).

51 Roy, V. et al. AI-2 analogs and antibiotics: a synergistic approach to reduce bacterial biofilms. Appl Microbiol Biotechnol 97, 2627–2638, doi:10.1007/s00253-012-4404-6 (2013).

52 Boyle, K. E., Monaco, H.T., Deforet, M., Yan, J., Wang, Z., Rhee, K., Xavier, J.B. Metabolism and the Evolution of Social Behavior. Mol Biol Evol. 34, 2367–2379, doi:doi: 10.1093/molbev/msx174. (2017).

53 Letelier, J.-C., Cárdenas, M-L., Cornish-Bowden, A. From L’Homme Machine to metabolic closure: steps towards understanding life. J Theor Biol. 286, 100–113, doi:10.1016/j.jtbi.2011.06.033. (2011).

54 Noble, D. A theory of biological relativity: no privileged level of causation. Interface Focus 2, 55–64, doi: 10.1098/rsfs.2011.0067. (2012).

55 Cárdenas, M. L., Benomar, S. and Cornish-Bowden, A. Rosennean complexity and its relevance to ecology. Ecological Complexity 35, 13–24, doi:org/10.1016/j.ecocom.2017.04.005 (2018).

56 Traore, A. S., Fardeau, M. L., Hatchikian, C. E., Le Gall, J. & Belaich, J. P. Energetics of Growth of a Defined Mixed Culture of Desulfovibrio vulgaris and Methanosarcina barkeri: Interspecies Hydrogen Transfer in Batch and Continuous Cultures. Appl Environ Microbiol 46, 1152–1156 (1983).

57 Nölling, J. et al. Genome sequence and comparative analysis of the solvent-producing bacterium Clostridium acetobutylicum. J Bacteriol 183, 4823–4838, doi:10.1128/JB.183.16.4823-4838.2001 (2001).

58 Bender, K. S. et al. Analysis of a ferric uptake regulator (Fur) mutant of Desulfovibrio vulgaris Hildenborough. Appl Environ Microbiol 73, 5389–5400, doi:10.1128/AEM.00276-07 (2007).

59 Bassler, B. L., Wright, M., Showalter, R. E. & Silverman, M. R. Intercellular signaling in vibrio-harveyi - sequence and function of genes regulating expression of luminescence. Molecular Microbiology 9, 773–786, doi:10.1111/j.1365-2958.1993.tb01737.x (1993).

60 Neufeld, J., D., Vohra, J., Dumont, M, G., Lueders, T., Manefield, M., Friedrich, M, W., Murrell, J, C. DNA stable-isotope probing. Nature Protocols 2, 860–866 (2007).

61 Neufeld, J. D., Schäfer H, Cox, M.J, Boden, R, McDonald, I.R, Murrell, J.C. Stable-isotope probing implicates Methylophaga spp and novel Gammaproteobacteria in marine methanol and methylamine metabolism. ISME J 1, 480–491, doi:10.1038/ismej.2007.65 (2007).

62 De Keersmaecker, S. C., Varszegi, C., van Boxel, N., Habel, L.W., Metzger K, Daniels, R., Marchal, K., De Vos, D., Vanderleyden, J. Chemical synthesis of (S)-4,5-dihydroxy-2,3-pentanedione, a bacterial signal molecule precursor, and validation of its activity in Salmonella typhimurium. J Biol Chem. 280, 19563–19568, doi:doi:10.1074/jbc.M412660200 (2005).

63 Lowery, C. A., Park; J., Kaufmann; G.F., Janda, K.D. An unexpected switch in the modulation of AI-2-based quorum sensing discovered through synthetic 4,5-dihydroxy-2,3-pentanedione analogues. J Am Chem Soc. 130, 9200–9201 doi:10.1021/ja802353j (2008).

